# Variability in the effects of transcranial direct current stimulation on free choice behaviour

**DOI:** 10.1101/2024.08.23.609379

**Authors:** Brandon Caie, Gunnar Blohm

## Abstract

Transcranial direct current stimulation (tDCS) is used as a tool to causally influence neural activity in humans non-invasively. Although most studies recruit a large number of participants in order to uncover population-level effects, growing evidence suggests that tDCS may be expected to induce different effects in different individuals, leading to large inter-individual variability and confounds in population-level testing. Additionally, variability may arise from intra-individual differences and confounds that are difficult to assess in studies with limited to no re-testing. Here, we performed 10 sessions of tDCS each on 5 human participants performing a free choice saccade task while neural activity was measured via EEG. Participants first underwent functional MRI to localize the human right frontal eye field (rFEF) homologue. An HD-tDCS montage was then used to focally target rFEF based on individual MRI localizations, alternating the polarity between anodal or cathodal current over repeated sessions during a 5 week period (twice weekly). On stimulation days, participants performed a free choice task prior to and after administration of tDCS while EEG activity was recorded. To quantify the likelihood that tDCS induced a causal effect on behaviour and neural activity across different levels of analysis, we developed a multilevel causal inference method based on permutation testing of a quasi-experiment (difference-in-differences). We then used this method to assess the likelihood of a causal effect at different levels of abstraction: group-level, participant-level, and paired-session level. At the group-level, we found evidence for an influence of tDCS on choice reaction times, which followed a reaction-time dependent change in EEG activity, and on how choices depended on previous trials. However, individuals showed heterogeneous effects. Further, analysis of paired-session variability often belied the pooled individual effect, suggesting that different sessions of tDCS may have produced markedly different effects in the same participants. In light of this, we discuss potential causes of this variability, and the counterfactuals that should be considered when making data-driven inferences about the effects of tDCS.

**Author Summary:** Developing reliable interventions on human neural activity is important for establishing causal relationships in basic research and developing therapeutics for pathological brain states. Transcranial direct current stimulation is a promising technique to intervene on neural activity safely in humans, but it is poorly understood if tDCS reliably impacts brain and behaviour in the same way across sessions. We performed an extensive test-retest study on tDCS in humans, and developed statistical methods to assess variability across sessions. Our results provide strong evidence against a consistent effect of tDCS in the same individual across different sessions. This warrants caution in using tDCS as a predictable intervention on neural activity in research and clinical practices.

## Introduction

It has long been known that externally applied electric fields over the cortical surface can influence neural activity (Bindman et al., 1964; Purpura and McMurty, 1965). Transcranial direct current stimulation (tDCS) was subsequently developed as a method to causally influence neural activity non-invasively (Merton and Morton, 1980; Rothwell, 2018). tDCS has several advantages: it is non-invasive, cost-effective, well-tolerated and easy to use. Early use of tDCS in humans suggested that cortical activity could be predictably modulated according to the polarity of the applied current (Nitsche and Paulus, 2000, 2001; Pellicciari et al., 2013; Priori et al., 1998). While this finding generated much interest in the experimental and clinical applications of directly intervening on neural activity in humans, future studies using tDCS were marred by confounding sources of variability and failures to replicate findings (Horvath et al., 2014; B. Krause and Cohen Kadosh, 2014).

The use of tDCS to influence human neural activity was motivated by studies in animal models, where slice preparations or invasive in vivo measurements isolated mechanisms of action at a resolution inaccessible to human study (Bindman et al., 1964; Pelletier and Cicchetti, 2014; Purpura and McMurty, 1965). It is now generally supported that anodal stimulation can increase, while cathodal stimulation can decrease, neural firing rate through a net polarization along a somato-dendritic axis (Kabakov et al., 2012; Liu et al., 2018), although it is unlikely that this is the sole physiological mechanism at play in vivo (Lefaucheur and Wendling, 2019; Rahman et al., 2013; Stagg and Nitsche, 2011). Thus, studies using tDCS were often motivated with this mechanism of action as an *a priori* that could be used to dissociate experimental or clinical hypotheses (Jackson et al., 2016; Lefaucheur and Wendling, 2019). However, computational models of current flow have shown that individual anatomical differences – such as skull thickness and composition, CSF volume, and gyri-sulcal morphology – can lead to different electric field distributions across individuals when the ‘same’ brain region is targeted (Bikson et al., 2012; Datta et al., 2012; Mikkonen et al., 2020; Opitz et al., 2015; Rawji et al., 2018; Truong et al., 2013). Given these known sources of variability, even if tDCS could be reduced to a singular mechanism of action (sometimes called the ‘somatic doctrine (Jackson et al., 2016)), predictions about the physiological outcome in absence of detailed knowledge of how individual neuroanatomy relates to the stimulation protocol may be expected to yield mixed and/or contradictory effects at the population level (Filmer et al., 2019; Vergallito et al., 2022).

With the recognition that tDCS may not yield consistent effects across individuals comes the need for causal assessments at the individual level. However, this introduces new methodological challenges. For one, individualized stimulation protocols that predict current distributions across the cortical tissue require imaging technology that negates the cost-effectiveness of tDCS. But perhaps more fundamental to the experimentalist, where effects are often *inferred* through statistical changes to behaviour and/or neural measurements, is the necessity of individual test-retesting – this requires significantly greater time investment on behalf of the participant, and also introduces statistical dependencies that can be abstracted over when testing hypotheses at the population-level (Caie and Blohm, 2024; Molenaar, 2004; Molenaar and Campbell, 2009). Knowledge concerning the variability of responses to tDCS within individuals is sparse, and studies assessing this variability typically rely on limited samples that would not be considered suitable for determining a population-level effect (Hsu et al., 2016; Willmot et al., 2024). Thus, it is often unclear to what degree the marked variability of tDCS responses can be attributed to functional differences in the mechanism of action of tDCS itself on different individuals, or to sources of variability that may arise either mechanistically (Ridding and Ziemann, 2010; Vergallito et al., 2022, 2023) or inferentially (Caie and Blohm, 2024; Granger and Newbold, 1974; Macke and Nienborg, 2019)) within individuals.

Here, we performed an extensive test-retesting study of tDCS in human participants, and developed a multilevel causal inference method to assess the likelihood of a tDCS-induced effect at different levels of abstraction: group-level, individual-level, and session-level. Individual fMRI localizations of the human homologue of the right frontal eye field (rFEF) were performed prior to the delivery of twice-weekly alternating sessions of anodal or cathodal current using high-definition transcranial direct current stimulation (HD-tDCS) for 5 participants over a 5 week period. Prior to and following administration of HD-tDCS, participants performed a free choice saccade task (Caie et al., 2023; Noudoost and Moore, 2011; Soltani et al., 2013) while EEG was recorded. Using our multilevel approach, we assessed the likelihood that pre-post differences to psychometric measures and EEG recordings were influenced by the polarity of the stimulation at different levels of abstraction: group-level, individual-level, and session-level. At each level, we found evidence for causal effects of tDCS on choice psychometrics, and corresponding changes in EEG activity. However, at both the group and inter-individual level, the variance in its constituent repeated measures was large and often contradictory. Indeed, an analysis of session-session variability supported the conclusion that different sessions of the same stimulation could produce opposite effects. We argue that this is sufficient evidence to call into question the consistency of tDCS effects within and across individuals. We then discuss the counterfactuals that must be evaluated in order to make inferences about the effects of tDCS through observation of its joint effects on neural activity and behaviour.

## Results

The study was composed of 5 participants, who underwent 10 sessions of fMRI-guided HD-tDCS, alternating between anodal and cathodal stimulation over an approximately 5 week period. Prior to and following stimulation, a free choice task was performed while EEG activity over the right frontal eye field (rFEF) was recorded. Thus, we aimed to determine whether the polarity of the stimulation changed how behaviour and/or EEG activity differed between pre and post-stimulation. By recording multiple sessions from the same participants, we were able to evaluate this at different levels of abstraction: Group, inter-individual, and intra-individual. Fig. 1 illustrates the relationship between higher and lower levels of abstraction, i.e. higher-level relationships are constructed by deriving summary statistics, such as averages, of the constituent relationships of a lower-level. At the group-level, studies aim to collect enough participants such that an average response within a sample generalizes to the average response of a population. The inter-individual level of analysis in turn asks whether group-level averages generalize to individual participants, while intra-individual analysis asks if the average effects on an individual are a consistent and representative measure of across-session variability. Further, the causal effect may be hypothesized to influence certain categories of trials in different ways (i.e. slowing decision-making following leftward choices and accelerating it following rightward choices).

**Figure 1:**
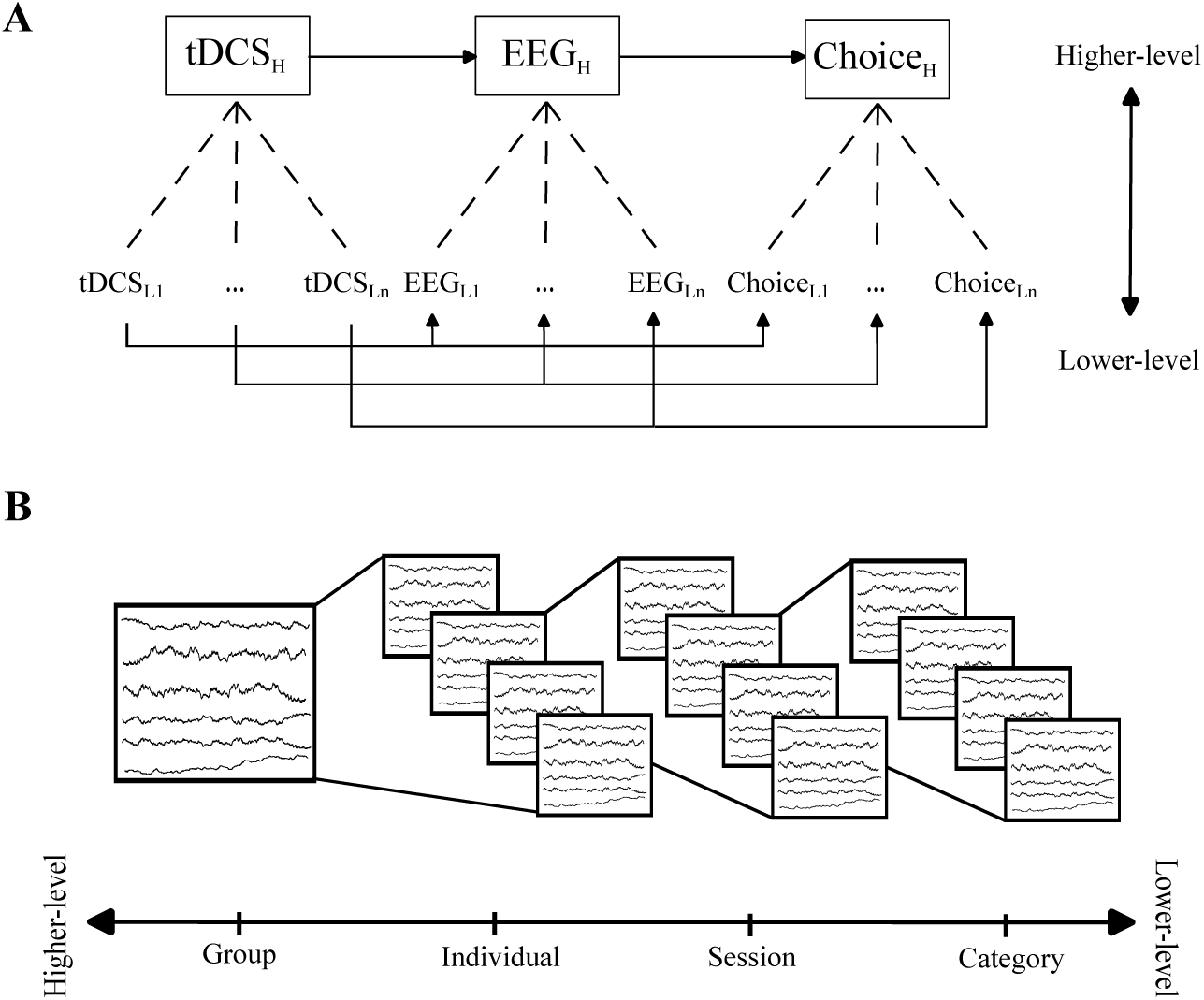
Levels of Abstraction. **A:** Causal diagram depicting the influence of tDCS on EEG activity, and EEG activity at different levels of abstraction. The validity of higher-level abstractions relies on the statistical independence of repeated measures of lower-level abstraction. **B:** Levels of abstraction analyzed in this paper. Group-level analysis are composed of repeated measures of individual participants, in turn are composed of repeated measures of individual sessions, are in turn composed of different ways we categorize different trials.

### Free Choice Task

Participants performed a free choice saccade task prior to and after the administration of HD-tDCS targeting rFEF (Caie et al., 2023; Noudoost and Moore, 2011; Soltani et al., 2013). Each trial consisted of a delay interval, followed by two targets presented asynchronously to each visual hemifield (Fig. 2). Participants directed a saccade to either target, as fast as possible, without anticipating. The temporal onset asynchrony (TOA) of the targets was randomized such that the preceding trial carried no additional information about the current trial. We quantified choice behaviour for each trial by recording the outcome of the choice (left or right) with respect to the TOA, that is the difference in time between the two target onsets, and the reaction time of the saccade relative to the onset of the first choice target. We then fit psychometric curves to the choice direction probability as a function of the TOA (Fig. 2C). From this, we derived two measures that were used to parameterize how choice direction depended on the TOA: the point of equal selection (*PES*), or the TOA at which there was an equal selection probability between left and right choices, and the slope of the psychometric function *W*, which denoted how fast the psychometric function rose between the 25th and 75th right choice probability percentile. These two parameters were used as aggregate measures of choice bias and sensitivity respectively; the PES denotes a directional bias in the data, while the slope denotes how much a change in the TOA around the midpoint corresponds with a change in the choice probability. Psychometric functions were fit with psignifit in python (*Methods: Psychometric Analysis*). The group-average psychometric function indicates that average choice probability for the rightward target reliably increased as a function of the TOA, thus the TOA influenced the choice probability.

**Figure 2:**
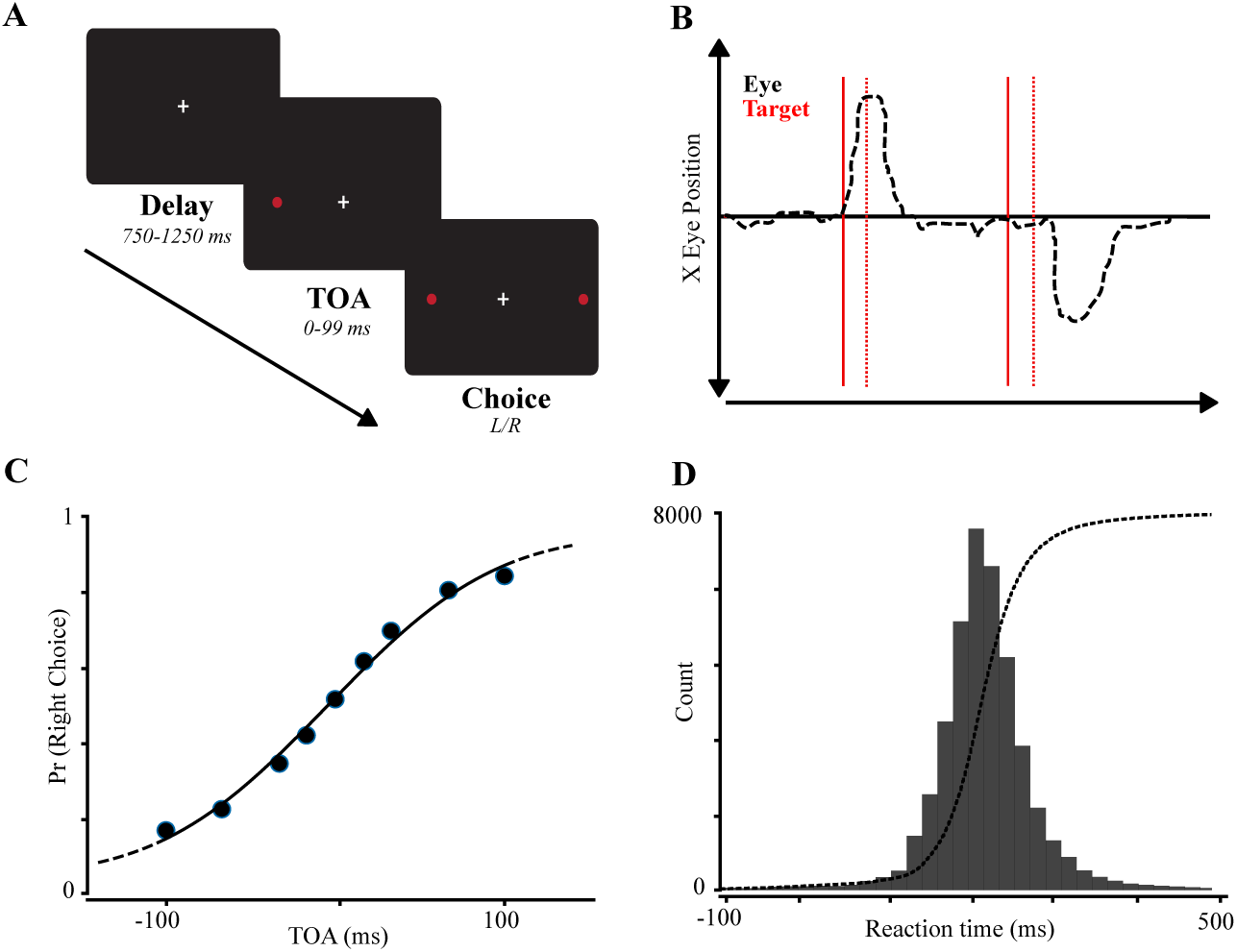
Free Choice Task. **A:** Free choice task schematic. Each trial begins with a randomized delay period, followed by the appearance of two choice targets. The temporal onset asynchrony (TOA) denotes the delay between the onset of the left and right target, with negative asynchrony values denoting the precedence of the left target. **B:** Eye trace of two trials. Positive horizontal eye position is right of the fixation cross. Slid red lines indicate leftward target, dashed red lines indicate rightward target. **C:** Group-average choice probability. Proportion of right target choices are plotted against the temporal onset asynchrony. Circles indicated average choice probability for a given TOA. **D** Histogram (grey bars) and cumulative response function (dashed line) of grouped reaction times. The horizontal red line indicates the appearance of the first target.

We also measured the reaction time of each trial, that is how long it took from the target onset to trigger a saccade. Models of the decision-process predict that reaction times are determined by the time it takes to integrate or amplify sequentially-sampled sensory information from an initial baseline evidence level to an action-triggering threshold (Carpenter, 1999; Carpenter and Williams, 1995; Cisek et al., 2009; Ratcliff and McKoon, 2008). In a bounded-accumulator framework, the mean reaction times (*µ_RT_*) directly correspond to the rate at which evidence is amplified or accumulated – therefore, we used mean reaction times as a proxy measurement for a component of the choice process that may not be directly accessible from measurements of choice probability. Additionally, we recorded the standard deviation of the reaction times (*σ_RT_*) as a proxy measurement for components of the choice process that may not change the mean reaction time, such as anticipatory responses, baseline evidence updating, and sensory noise (Caie et al., 2023; Kim et al., 2017; Nakahara et al., 2006). We stress that these are merely proxy measures for a model-based abstraction of the choice process that may have been influenced by tDCS, and are not intended to represent anything akin to a cognitive natural kind. Fig. 2D shows the empirical histograms (grey bars) and overlapped cumulative distribution (dashed lines) for the combined data. The same choice psychometrics and reaction time plots for the individual participants are then shown in Fig. 3.

**Figure 3:**
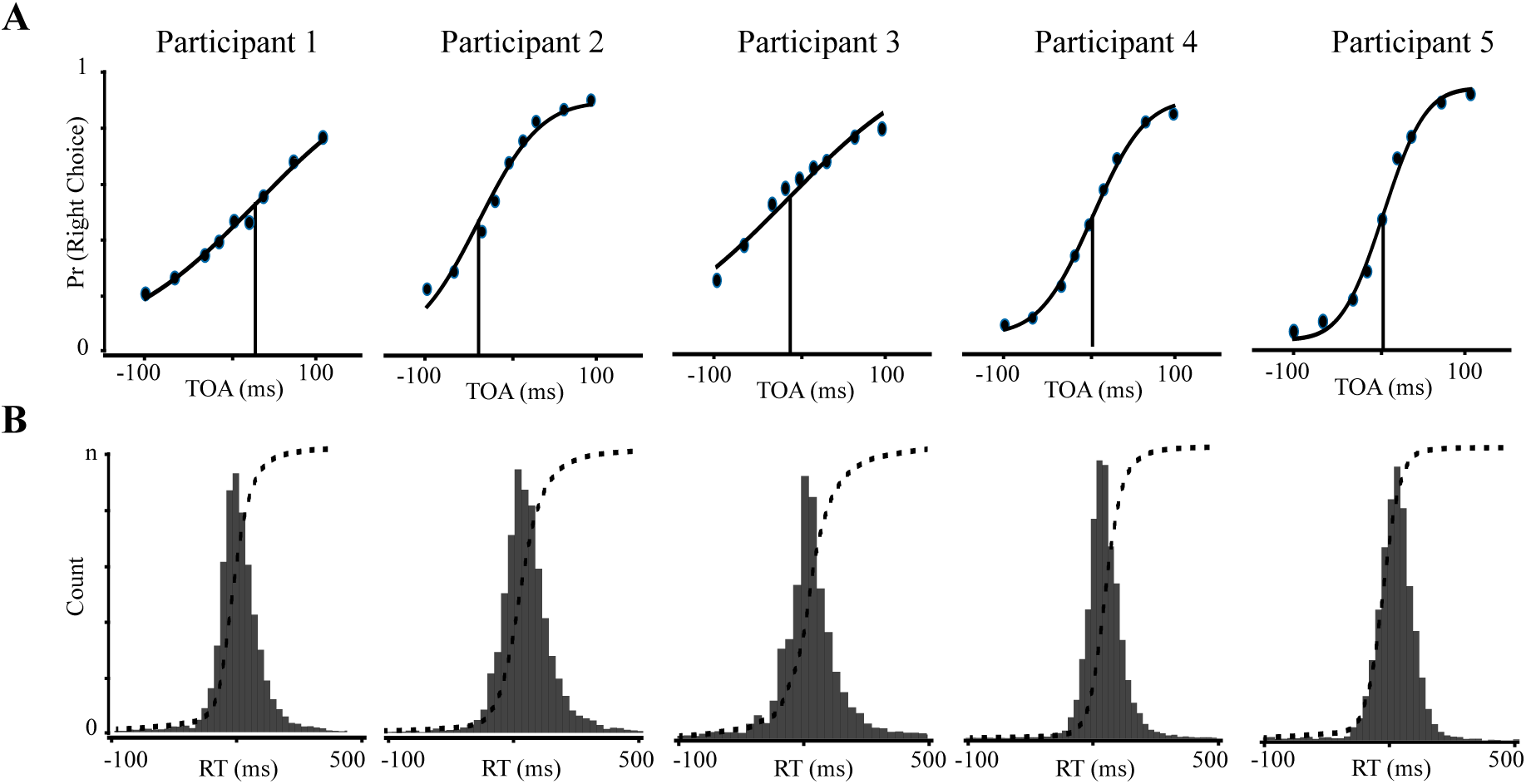
Individual Participant Psychometrics. **A:** Individual choice probabilities are plotted for the 5 participants. **B:** Reaction time histograms and cumulative response functions. Conventions as before in Fig 2.

### Permutation testing of difference-in-differences

We sought to assess the likelihood of a causal effect of tDCS at different levels of abstraction: group-level, individual-level, and session-level. Each of these levels implies different possibilities about the causal relationship between tDCS and neural activity. At the group-level, a positive finding may provide evidence for the likelihood of an average effect on a larger population. Although our study is not intended to be powered for providing this sort of analysis, as only 5 participants performed the study, we assess group-level findings in this manner so as to better illustrate the relationship between individual and session-level findings. At the individual-level, an effect may provide evidence for a causal effect of tDCS that is person-specific. Finally, a session-level analysis may provide evidence for a causal influence of tDCS that was specific to the context of that particular day.

To assess these different questions under one framework, we developed a multilevel causal inference method based on permutation testing of difference-in-differences (Methods: Multilevel approach to causality testing). Briefly, this method estimates the likelihood of a causal effect by computing a distribution of possible ‘quasi-experiments’ in which there was no difference between anodal and cathodal HD-tDCS. The method of difference-in-differences is a quasi-experimental approach designed for situations in which randomization is difficult, such as in many hypotheses tested in the social sciences (Marinescu et al., 2018. Although such an approach may not be necessary for population-level testing of tDCS, where randomization of anodal and cathodal stimulation would allow for confounding variables changing across pre and post-stimulation to be factored out, it is important for assessing variability that may occur across sessions, where each session can (and should) be considered its own quasi-experiment with a potentially different effect. The procedure is depicted in Fig. 4. Thus, instead of treating pre and post-stimulation blocks as independent events and assessing their difference, we treat them as sequential processes where an effect may be confounded by underlying non-stationarities. We then test whether the ‘difference-in-difference’, that is the relative difference between how anodal and cathodal stimulation influenced the difference between pre and post-stimulation blocks, was itself different than would be expected by chance from random permutations of the data.

**Figure 4:**
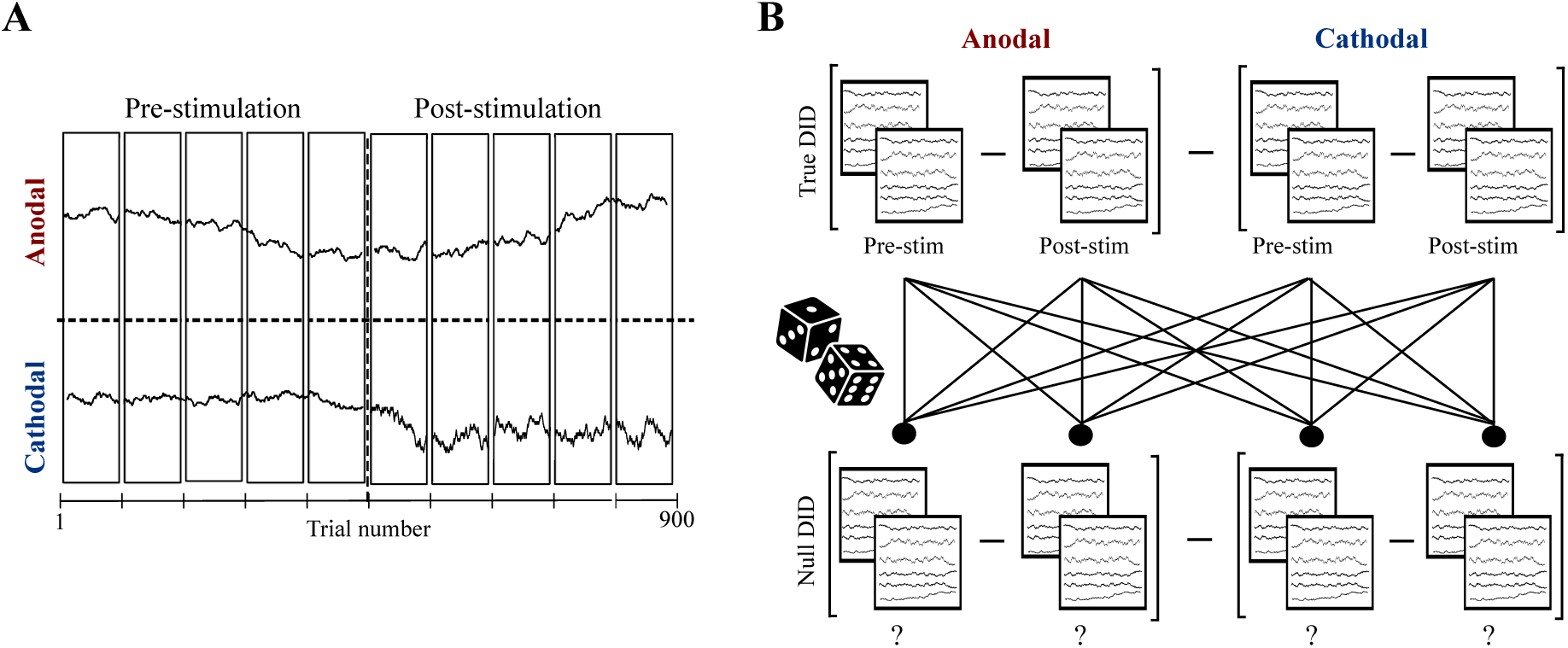
Permutation Testing of Difference-In-Differences. **A:** Time series representation of a single tDCS session. Depicted is an arbitrary time series for anodal (top) and cathodal (bottom) sessions, demarcated by the dashed horizontal line. Pre and post-stimulation epochs are demarcated by the vertical horizontal line. Sessions were organized into 5 blocks of 90 trials for both pre and post-stimulation. Pre and post stimulation were separated by a 20 minute interval (not shown). **B:** Cartoon depiction of permutation testing, Top: for a given metric, the true difference-in-difference (DID) subtraction is obtained by subtracting the across-trial means from pre and post-stimulation for anodal (top left), subtracting the across-trial means from pre and post-stimulation for cathodal (top right), and then subtracting the difference of the two differences. The null DID distribution (bottom) is then obtained by randomly assigning trial indices to 4 evenly sized groups, performing the same subtraction, and iterating over a chosen number of permutations.

### Group-level testing

We first analyzed the full dataset by combining trials across all participants into separate groups comprising the pre and post-stimulation periods for anodal and cathodal sessions separately. At this level of analysis, no distinction between participants are made. Although this method of combining data has obvious interpretational issues for any group-level conclusions drawn from a dataset with 5 participants (despite the repeated measures within participants), we perform this analysis not to support the generalizability of its result so much as to assess how they correspond to within-participants variability, whatever the result may be. In doing so, we risk introducing a straw man for how group-level studies are conducted in the first place. In a sense this is true, however we contend that although group-level variability is not always combined across participants in such a blind fashion, data *within-participants* is.

Permutation testing on the group-averaged data was performed to estimate the null distribution of four psychometric parameters (the point of subjective equality (*PSE*), the sensitivity *W*, the median RT *mu_RT_*, and the RT variance *σ_RT_*) and on each point in the EEG time-frequency response matrix during the delay interval. Fig. 5 shows the outcome of permutation testing for the psychometric (Fig. 5A) and EEG time-frequency responses (*Fig 5B*) when all sessions and trials were combined across all participants. Fig. 5A) shows the empirical null distributions for the *PES* (top left), *W* (top right), *µ_RT_* (bottom left) and *σ_RT_* (bottom right) in grey, with the true difference-in-difference denoted by the solid vertical black line. Here, it can be seen that it was statistically likely that the difference-in-difference for both metrics used to quantify choice probability, (*PES* and *W*, could have arisen from random permutations of trials (*PES p = .264*, *W p = .392*). Thus, there is no evidence to suggest that the choice outcome was influenced by the polarity of the stimulation when all data was combined. However, both metrics for reaction times were statistically unlikely to have arisen from random permutations of trials (*µ_RT_* p < .001, *σ_RT_* < .001), suggesting that the polarity of stimulation was required to explain how reaction times differed between the pre- and post-stimulation period.

**Figure 5:**
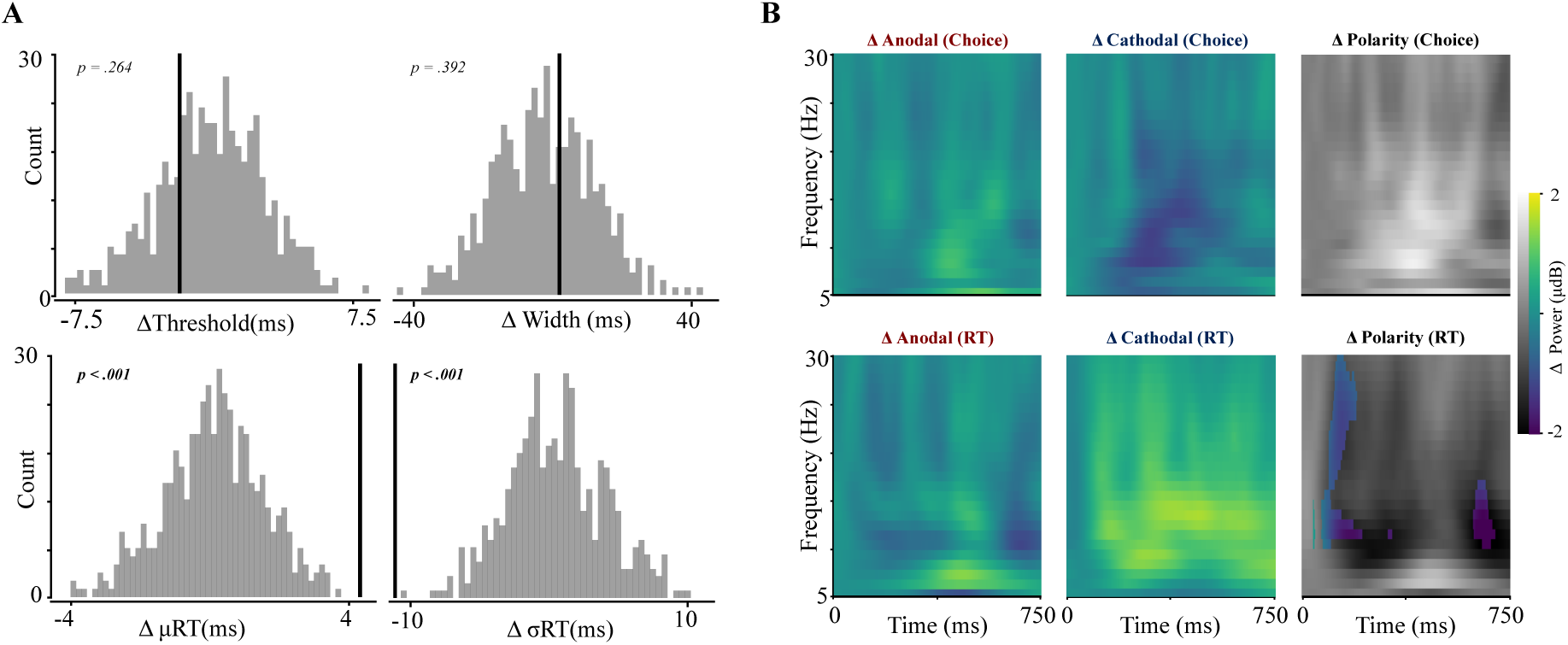
Group Permutation Testing. **A:** True DID (solid black bar) and empirical null distribution (grey bars) are plotted for the PSE (top left), width (top right), mean reaction time (bottom left) and RT standard deviation (bottom right). Significant p values are reported in bold. **B:** EEG time-frequency subtractions for left vs right choice (top) and slow vs fast RT (bottom). Depicted are the mean power subtractions (pre-post) for anodal (left) and cathodal (middle) sessions, and the polarity DID subtraction (right).

We then modified the difference-in-difference calculations in the psychometrics (where a single metric was chosen to perform a double subtraction on) to assess whether a change in behaviour corresponded with a change in an EEG time-frequency response that correlated with behaviour. To do this, we computed a triple-difference estimator (Olden and Møen, 2022) by including a median split between a given type of behavioural trial as an additional way to divide the data. We split behavioural trials in three ways here based on the outcome of the trial following the EEG activity: left vs right choice and a median split between slow and fast RT . For a given split, we had a difference-in-difference for each time-frequency point for anodal sessions that first subtracted the median split in pre-stimulation, then the median split in post-stimulation, and then subtracted these two differences to yield a single difference-in-difference for the anodal sessions. This was then performed again for the cathodal sessions, and the final triple-difference estimator was calculated as the subtraction between the two. The triple-difference was then compared statistically to an empirical null distribution that was calculated as before, where instead of dividing the data in permutations of 4 groups it is now 8.

Fig. 5B shows the EEG time-frequency triple-difference subtractions with respect to the upcoming choice direction (top row) and reaction time (bottom row) for the two session conditions (pre-post) for anodal (left) and cathodal (middle) sessions. We then computed the triple-difference subtraction, plotted in the right column. For the triple-difference subtraction, we plotted the true difference first in grayscale, with statistically significant time-frequency pixels overlayed in colour (sidak corrected for multiple comparisons, p values < .0001 reported). Here, it can be seen that, just as with the behavioural psychometrics, there was no evidence for a polarity-dependent effect on how the EEG time-frequency response changed with respect to the future choice direction. However, this subtraction performed relative to the upcoming reaction time revealed a significant relationship in alpha and beta-band activity in the early portion of the trial, and alpha-band activity in the late portion of the trial. Therefore, when combining data from all participants and all sessions together as a single functional unit of analysis, we found evidence for a behavioural effect of tDCS on reaction times that followed a change in the recorded EEG response.

Free choice saccadic behaviour has been demonstrated to be history-dependent (Caie et al., 2023; Soltani et al., 2013). It is thus conceivable that choice history-dependent effects of tDCS may be averaged together, and thus obscured, in trial-averaged analyses. For example, if anodal and cathodal stimulation differentially influenced the choice process depending on whether the previous choice was in the left or right direction, combining these different trials together could obscure the tDCS-dependence in the case that these two types of trials were balanced. We therefore assessed whether the influence of tDCS on choice psychometrics and EEG time-frequency responses was dependent on information from the previous trial (*Methods: Permutation Testing*) We used three different metrics for the previous trial: the direction of the previous choice (a spatially-dependent history effect), whether the current choice corresponded to a repetition or an alternation of the previous choice (a spatially-independent history effect), and whether the previous trial resulted in a slow or a fast RT relative to the median. These outcome measures reflect three distinct abstractions of feedback that may occur following a trial: the previous direction split indexes whether tDCS influenced the trial-trial buildup of a directional bias (Dorris et al., n.d.; Fecteau et al., 2004, the repetition split indexes a trial-trial buildup of sequence effects that may not be specifically lateralized (Cho and Cohen, 2002; Gao et al., 2009), and the reaction time split indexes a metric of task performance analogous to post-error slowing (Dutilh et al., 2012; Laming, 1979), given the requirement of participants to choose as fast as possible.

We compared the true triple-difference estimator to the permuted null distribution for these three outcome measures. Fig. 6 shows the results for the repetition/alternation difference (top row), previous choice difference (middle row) and previous reaction time difference (bottom row). With respect to the psychometrics, we found evidence for an effect of the previous choice direction on the PES (*p < .001*), and an effect of the previous reaction time on the *W* (*p = .016*) and *σ_RT_* (*p = .05*). In contrast to the group-averaged choice and reaction time subtractions, statistical testing of the EEG time-frequency responses did not perfectly correspond with the psychometrics. While the direction of the previous choice was a better predictor of the influence of polarity on psychometrics, the identity of the previous choice *relative to the current choice* (repetition vs alternation) was a better predictor of changes to the EEG time frequency response, with a modulation in both the alpha and beta band surviving statistical significance. However, we did observe a modulation of beta-band activity with respect to the previous reaction time that co-occurred with the change to the width and RT variance. Taken together, we observe that the outcome of the previous trial was predictive of how choice psychometrics and EEG activity depended on tDCS polarity, however only when conditioning on the previous reaction time did we find a correspondence between statistical changes to the psychometrics and the EEG.

**Figure 6:**
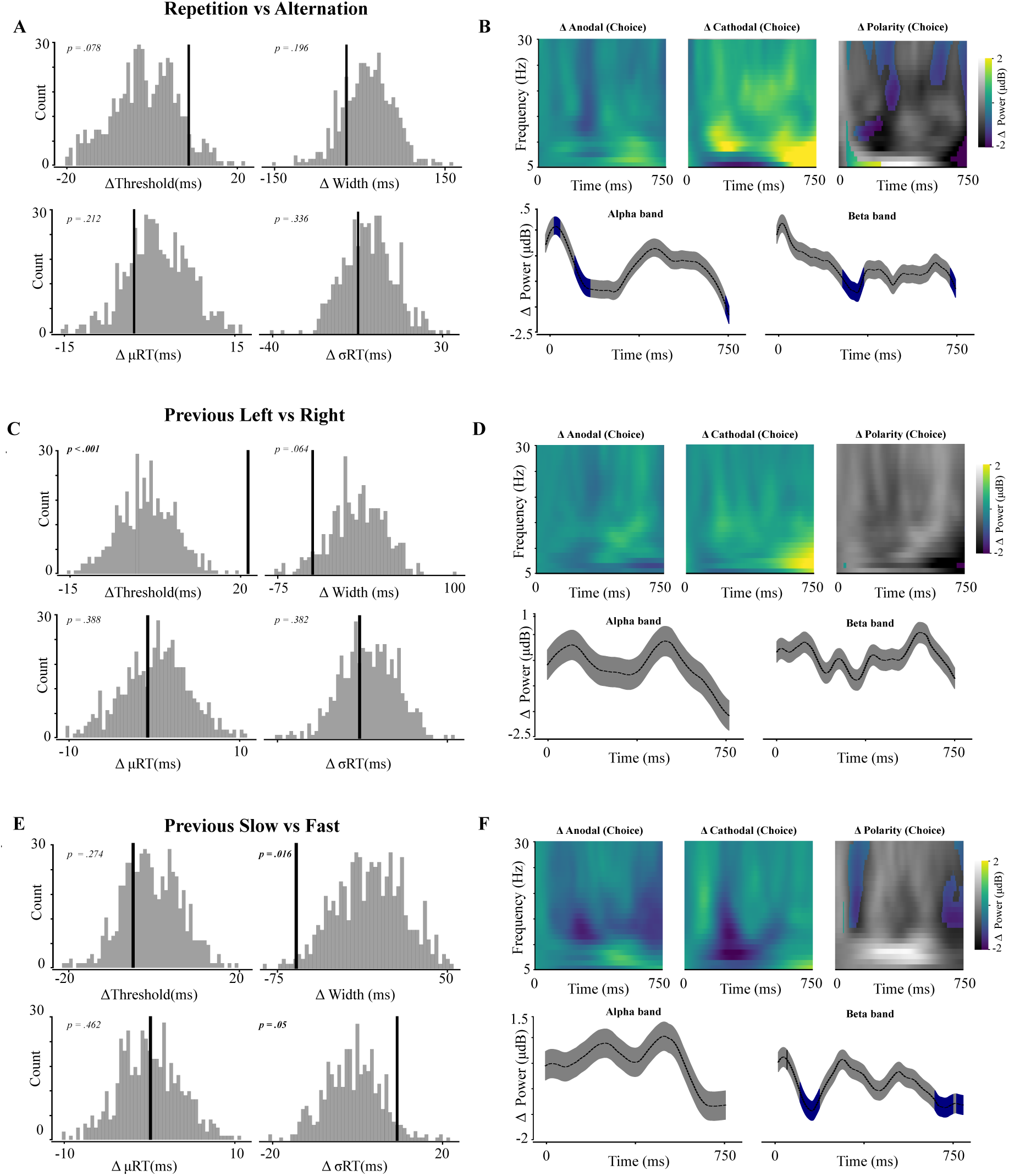
Choice History Permutation Testing. **A-B:** Permutation testing for triple difference of repetition vs alternation trials across all participants. **A:** depicts the psychometric true TD and null distribution as in Fig 6. **B:** EEG time-frequency response triple-difference. Shown are the repetition/alternation difference-in-difference (DID) for anodal (top left) and cathodal (top middle), from which the polarity TD is computed (top right). Significant TFR pixels are in colour. Bottom left: Time-series of alpha band (7Hz band, mean power triple-difference in black, std power triple-difference in grey, significant time-points shaded in blue). Bottom right: The same, for the beta band (15 Hz). **C-D:** Previous left vs right choice triple difference. Plots are otherwise the same. **E-F:** Previous reaction time (fast vs slow median split)

### Inter-individual testing

Group-level analyses of tDCS studies have previously been shown to be comprised of largely heterogeneous results at the inter-individual level. This may result from fixed anatomical-physiological differences that result in variable effects being averaged together at the group level, or unexplained variability within participants that is erroneously interpreted as relevant to a causal effect of tDCS. Previously, it has been shown that the induced electric field for a given target site can vary dramatically across participants depending on individual morphological differences. Our stimulation protocol (*Methods: fMRI-guided HD-tDCS*) was designed so as to minimize confounding variables by a) minimizing current spread with a 4×1 HD-tDCS electode montage (cite sources on that) and b) selecting a region of interest that corresponded to the individually-specific neural activation close to the putative human homologue of the right frontal eye field (frontal eye field where art thou), as measured through an fMRI saccade localizer task.

The resulting electrode montages for each participant are overlayed on individual 3D models of the scalp in Fig. 7, with the induced electric fields for an anodal stimulation session (center electrode 2mA, surround electrodes -.5 mA) overlayed on both the surface of the scalp (top row) and the gray matter (bottom row). Visually, variation in the location, intensity, and spread of the current can be observed. We depicted the distribution of current intensity for a single participant as a histogram of the electric field normal vector (measured in V/m) for each triangle in the mesh composing the gray matter surface, with the average and standard deviation of each electric field normal shown at the target ROI for each participant in Fig. 7C. Here, it can be seen that the intensity of the current was different for each participant. This confirms that inter-individual anatomical differences should be considered as a source of variation in outcome metrics, however with the number of participants in this study little systematic can be said about how current intensity and spread relates to the variability in the inferred effect across individuals. Coordinates for the region on the gray-matter and pial surface for the center electrode, and the positions of the surround electrodes on the pial surface for each participant are shown in Table 1.

**Figure 7:**
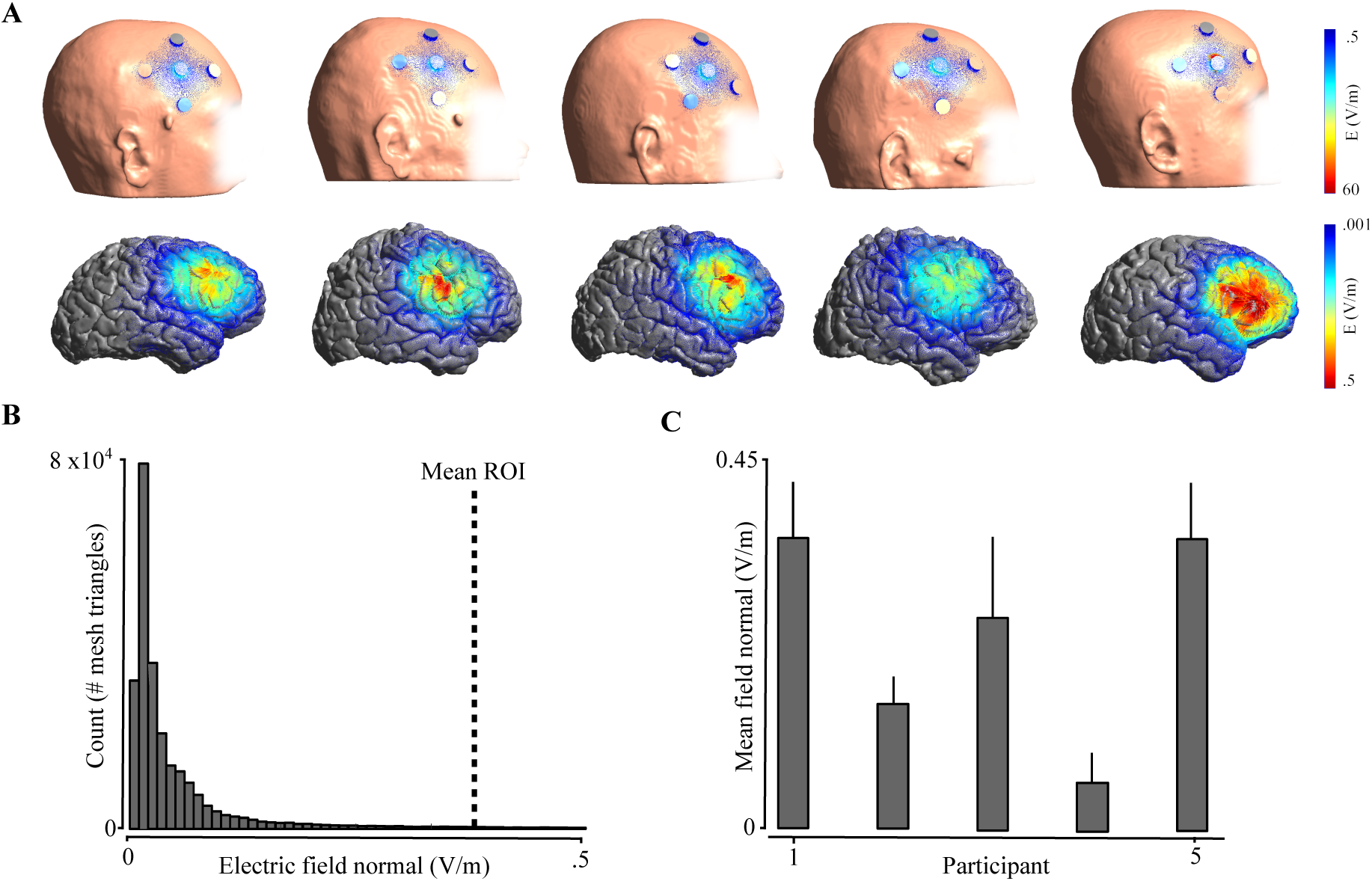
Individual current modelling **A:** Electric field simulations for each participant (left to right) are overlayed on the scalp (top row), showing the center-surround HD-tDCS ring setup, and the gray matter surface (bottom row). Electric fields are plotted in v/M. **B:** Distribution of normal component of electric field for an example participant. Histogram is shown for all triangles in the 3D mesh corresponding to the gray matter surface. **C:** Normal component of the electric field at the ROI for each participant. The mean and standard deviation are shown for mesh triangles corresponding to less than a 2cm radial distance from the ROI.

**Figure 8:**
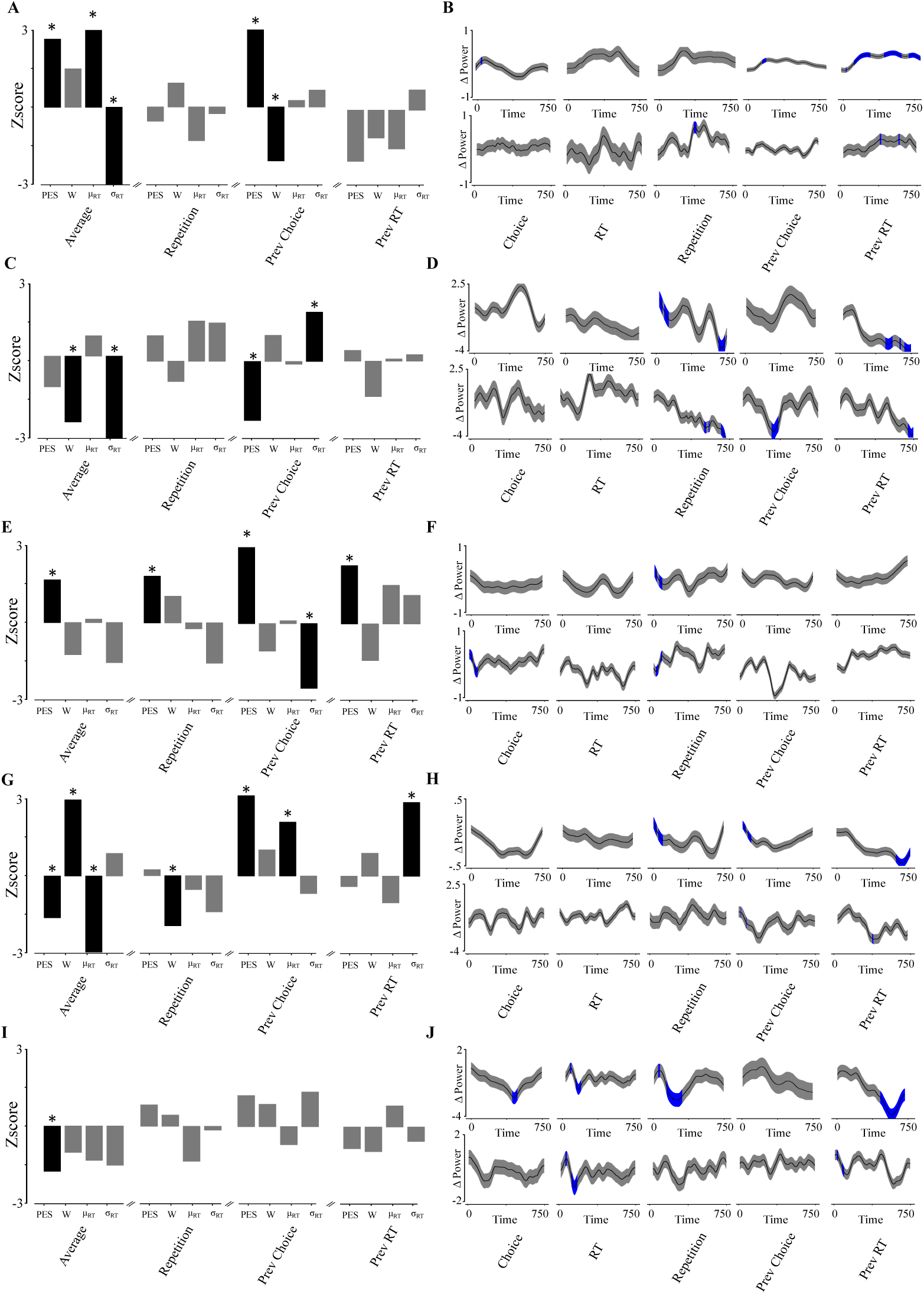
Inter-Individual Permutation Testing.**A-B:** Psychometric and EEG permutation testing for participant 1. **A:** Z-scores for the true DID relative to the permuted null distribution for PSE, W, *µ_RT_*, and *σ_RT_* . Significant tests are denoted by black bars and stars (left). TD permutation z-scores for repetition vs alternation (second from left), previous left vs right (second from right) and previous slow vs fast RT (right) are also reported. **B:** Top: mean alpha-band DID for choice subtraction, RT subtraction, and mean alpha-band TD for repetition, previous choice, and RT subtractions. Bottom: The same, for beta band. **C-D:** Participant 2. **E-F:** Participant 3. **G-H:** Participant 4. **I-J:** Participant 5.

**Table 1:**
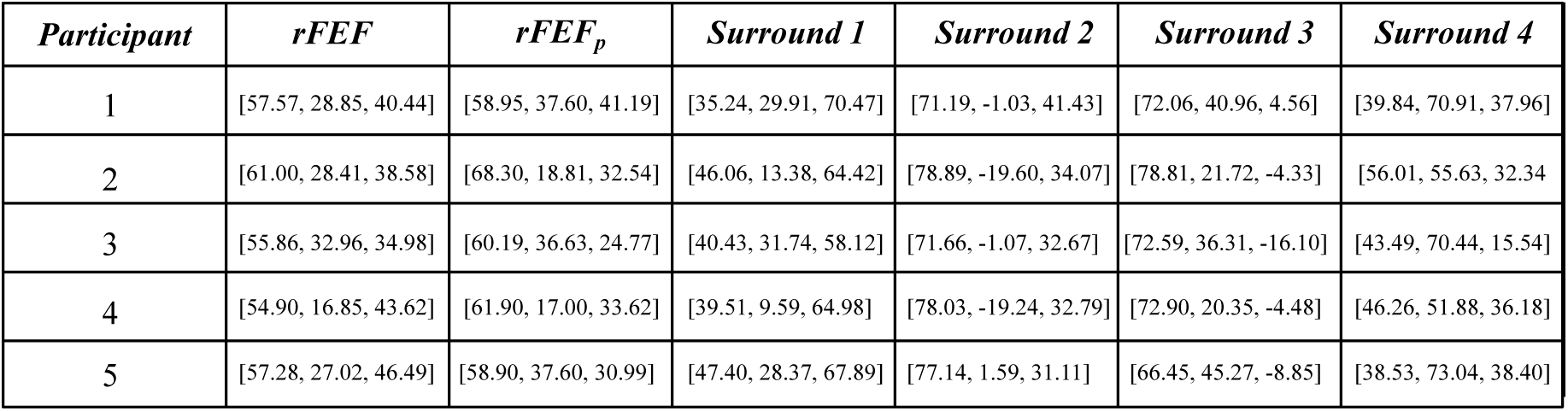
Table of ROI coordinates in SimNibs for each participant. *rFEF* corresponds to location of ROI projected onto the gray matter surface. *rFEF_p_* corresponds to the location of the center electrode, as projected onto the skin surface. Surround electrodes are also projected onto the skin surface.

| *Participant* | *rFEF* | *rFEF<sub>p</sub>* | *Surround 1* | *Surround 2* | *Surround 3* | *Surround 4* |
| --- | --- | --- | --- | --- | --- | --- |
| 1 | [57.57, 28.85, 40.44] | [58.95, 37.60, 41.19] | [35.24, 29.91, 70.47] | [71.19, -1.03, 41.43] | [72.06, 40.96, 4.56] | [39.84, 70.91, 37.96] |
| 2 | [61.00, 28.41, 38.58] | [68.30, 18.81, 32.54] | [46.06, 13.38, 64.42] | [78.89, -19.60, 34.07] | [78.81, 21.72, -4.33] | [56.01, 55.63, 32.34] |
| 3 | [55.86, 32.96, 34.98] | [60.19, 36.63, 24.77] | [40.43, 31.74, 58.12] | [71.66, -1.07, 32.67] | [72.59, 36.31, -16.10] | [43.49, 70.44, 15.54] |
| 4 | [54.90, 16.85, 43.62] | [61.90, 17.00, 33.62] | [39.51, 9.59, 64.98] | [78.03, -19.24, 32.79] | [72.90, 20.35, -4.48] | [46.26, 51.88, 36.18] |
| 5 | [57.28, 27.02, 46.49] | [58.90, 37.60, 30.99] | [47.40, 28.37, 67.89] | [77.14, 1.59, 31.11] | [66.45, 45.27, -8.85] | [38.53, 73.04, 38.40] |

We now describe the results of permutation testing for psychometrics and EEG time-frequency responses at the inter-individual level. Fig. 9 plots the z-score of each difference-in-difference for all trials (left), and the previous-trial triple-difference conditioned on repetitions vs. alternations (middle-left), previous left vs. right choice (middle-right), and previous slow vs. fast RT (right), with z-scores corresponding to a p-value below 0.05 in black. First we assessed whether the null effect of choice probability and positive effect of reaction times was recapitulated consistently at the individual level. Here we found that within individuals, contrary to the group-level findings, choice probability (as indicated by the *PES* and *W* parameters in the leftmost columns) was statistically distinguishable in all participants, however in contrasting directions (*PES subtraction:* increased in 2/5 participants, decreased in 2/5, null change in 1/5; *W subtraction:* increased in 1/5, decreased in 1/5, null in 3/5). Thus, the null effect observed at the group-level was composed of a mixture of null and contrasting positive results that averaged as a non-significant effect. In the case of reaction times, where we observed a group-level significance for both the mean and standard deviations, we again found hetereogeneity in the individual data. For the standard deviation, there was some consistency, however only two participants showed significant interactions with the tDCS polarity. For *µ_RT_*, again only two participants (1 and 4) showed significant changes, however now in opposite directions.

**Figure 9:**
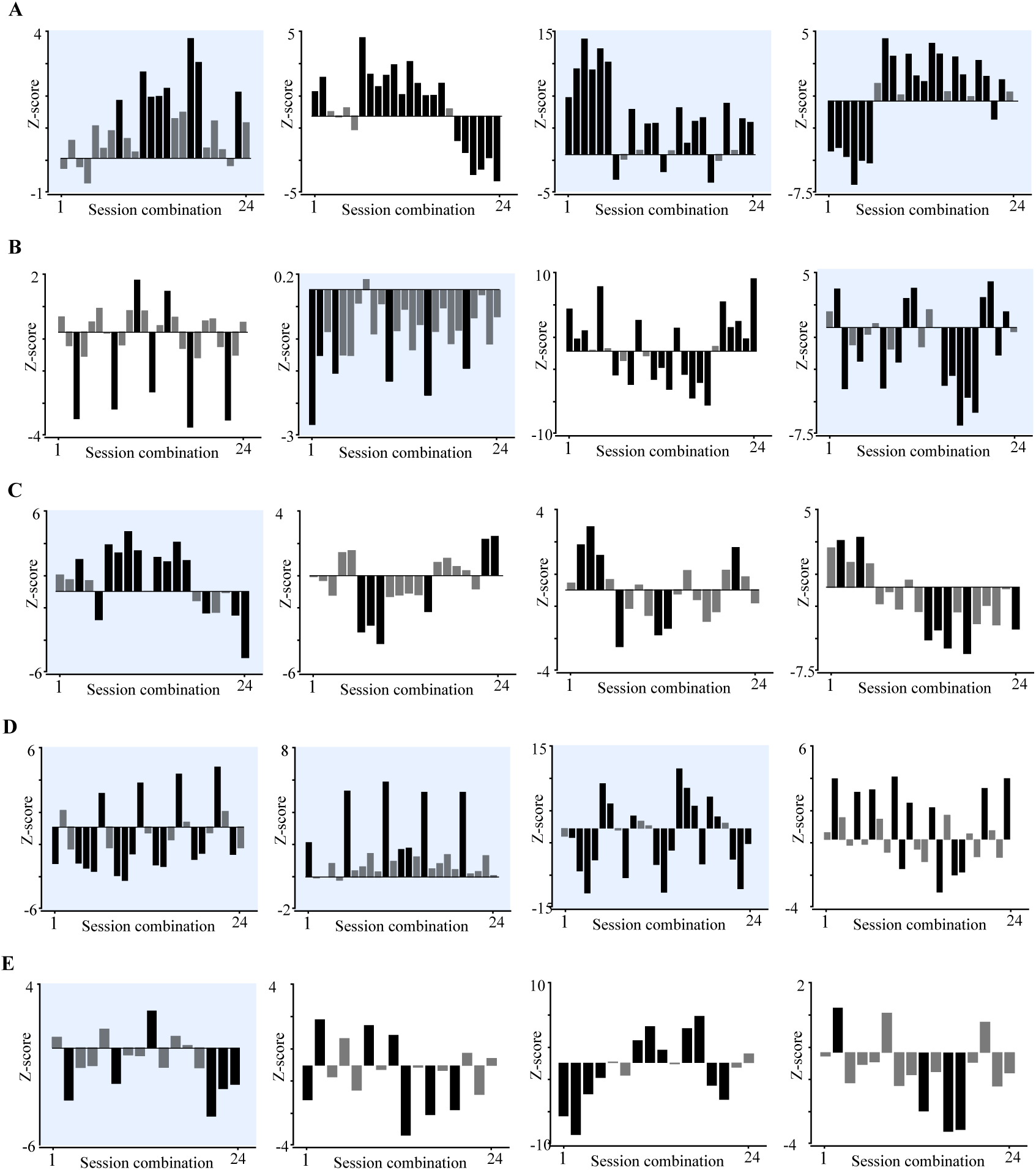
Intra-individual variability in psychometric subtractions at the session-pairing level. **A:** Participant 1 Z-scores for each possible session-pairing permutation are plotted for *PSE, W, µ_RT_, σ_RT_*, left to right. Black bars denote z-scores corresponding to a significant relationship (p<.05), grey bars denote p > .05. Plots shaded in blue denote where significance was reached for the corresponding psychometric subtraction when combining data across sessions. **B:** Participant 2. **C:** Participant 3. **D:** Participant 4. **E:** Participant 5

At the group-level, we did not observe a significant change in the choice-dependent EEG activity. However, within participants we found a significant change in the choice-dependent alpha-band activity in 2 of 5 participants, and a change in the choice-dependent beta-band activity in 1 of 5 participants. Fig. 9B plot the mean and standard deviation of the choice (left) and RT-dependent (middle left) alpha (top) and beta-band (bottom) power subtractions, with significant time-points (multiple comparison corrected p < .001) overlayed in blue. In contrast, at the group-level we did observe a reaction-time dependent modulation of alpha and beta-band activity, although a significant relationship at the individual participant level was only found in a single participant, suggesting that the detection of this effect resulted from an averaging effect that poorly explained the inter-individual variation. As such, we cannot conclude that the group-level statistics we reported previously are supported by a consistent relationship with inter-individual psychometrics. We note that this should not be unexpected, given our sample size.

We then performed the same analysis on choice history, using the same variables (repetition/alternation, previous choice direction, previous RT median split) as with the group-average. While the group-level averaging only resulted in a joint effect of history on psychometrics and EEG activity for the previous RT split, we report a diversity of effects reaching significance within participants here. For the repetition/alternation split, we see a change in the choice psychometrics for two participants, but alpha and beta-band changes surviving the significance threshold for all participants. Notably, these modulations occurred with distinct profiles across participants. Repetition-dependence on stimulation polarity resulted in a diversity of temporal profiles of alpha and beta modulation such that a clean distinction between an increase and decrease in band-power was difficult to assess – for example, in participant 2 (Fig.9B), there was an observed power difference in alpha activity that increased in the early portion of the trial, and decreased in the late portion.

In the previous choice split, we find additional evidence for heterogeneity in the psychometric effects, with 3 of 5 participants showing an effect of *PES* in one direction, and 1 participant showing an effect in the opposite direction. Similarly, 2 of 5 participants showed an effect of *σ_RT_*, again in opposing directions, with no corresponding consistency observable in either EEG bands analyzed here. Notably, in the previous RT split, having previously observed a significant group-level relationship in the *W* and *σ_RT_*, only a single participant survived significance testing for either of these metrics (participant 4, *σ_RT_ p* = *xx*. The group-level change in *W* appears to have been driven by individual participant effects that were in the same direction for 4/5 participants, but none surviving significance testing in isolation. For *σ_RT_*, the positive result seen in the group-level was composed of a single positive result at the group-level (participant 4), plus 3 participants with nonsignificant trends in this direction, and one participant with a nonsignificant trend in the other direction. As with the other findings, EEG alpha and beta-band activity was likewise heterogeneous. Although no group-level alpha-band relationship was observed, an individual alpha-band relationship was found in 2 of 5 participants, however in opposite directions with respect to changes in band-power. Conversely, a group-level beta-band relationship was observed, but again we found a mixture of previous RT-dependent increases and decreases across participants.

### Intra-Individual

The previous analyses of inter-individual psychometrics and EEG time-frequency responses aggregated all of the tDCS sessions together, and split them by the polarity of the stimulation on that day only. We now appraise these findings in light of the variability observed across-sessions. Our method of assessing intra-individual variability was to treat every possible anodal and cathodal session pairing within our dataset as a possible permutation that could be used in isolation to provide evidence for the underlying relationship of tDCS polarity at the individual level. In our permutation analysis, we preserved the order of the polarities such that the subtraction logic would be consistent by only including the session permutations with the anodal stimulation session as the first entry (*Methods: Session Permutations*). For a participant with 10 sessions, 5 anodal and 5 cathodal, this resulted in 25 possible session pairings that could be used to contrast the effects of anodal and cathodal tDCS. Our general analysis strategy was to see if previously significant findings at the inter-individual level corresponded to consistent relationships across different session pairings, and if this relationship between the psychometric subtractions at the individual session level predicted differences in the EEG response.

Fig. 10 shows the permutation z-scores for each session pairing within participants, plotted along rows, where the columns are separated by the choice psychometric (left: *PSE*, middle left: *W*, middle right: *µ_RT_*, right: *σ_RT_*). Session pairings that survived statistical significance (p < .05) are indicated with black bars, whereas session pairings that did not are indicated with grey. For comparative purposes, psychometric subtractions that survived significance testing at the inter-individual level are shaded in blue. The first observation that should be stated is that for every possible participant-psychometric combination, at least one possible pairing of anodal and cathodal sessions would have resulted in a statistically significant difference. This in itself is notable, because it calls in to question the notion that variability seen in inter-individual tDCS studies across participants reflects a veritable inter-individual difference, rather than a sample of a larger variability space than is measurable within the contexts of single paired-session studies. Naturally, such questions are specific to the variability imposed by the combination of the task and behaviour, the temporal sequence of the paired-sessions, and the sources of variability the participant imposes; because of this, these conclusions do not provide a refutation of any specific paired-session design *per se*, but offers a singular example where caution in the evidential value of single paired-sessions should be exercised.

**Figure 10:**
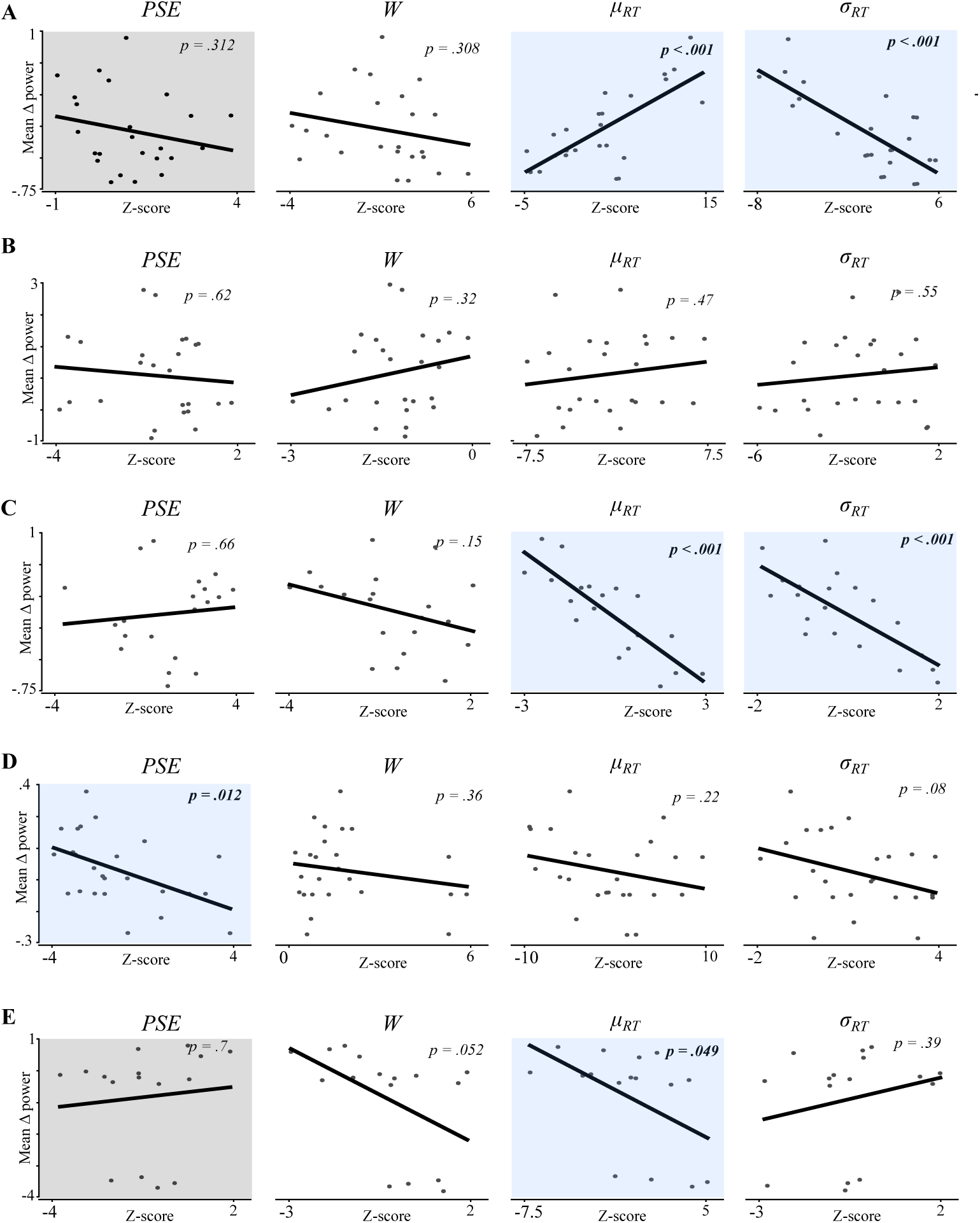
Session-wise correlations between psychometrics and EEG alpha-band differences. **A:** Double-subtraction z-scores are plotted against mean alpha-band double subtractions (averaged across trial timepoints) for *PSE, W, µ_RT_, σ_RT_* for participant 1. Grey boxes denote correlations where there was significance in both the psychometrics and EEG measures at the session-average level. Blue boxes denote significant session-wise correlations. **B:** Participant 2. **C:** Participant 3. **D:** Participant 4. **E:** Participant 5.

The second observation that should be noted is a disconnect between psychometric subtractions that survived significance at the inter-individual level, and the consistency of the session pairings that constitute it. While some psychometric subtractions that survived inter-individual significance testing were remarkably consistent across session-pairings (see Participant 1 *PSE, µ_RT_*, Participant 2 *W*, Participant 4 *W*), other psychometric subtractions surviving significance were composed of sessions that, when paired together, showed highly variable effects (see Participant 1 *σ_RT_*, Participant 2 *σ_RT_*, Participant 4 *PSE, µ_RT_*). Thus, it can be seen that for psychometric effects detected by combining repeated measures of anodal and cathodal sessions together, individual combinations of these sessions may yield strong effects in either the detected direction, or its opposite. Taken together, these assessments could be used to suggest that only psychometric subtractions with a consistent relationship at the session-level should be viewed as evidence, due to averaging confounds arising from the likelihood of detecting effects at the single paired-session level. Alternatively, we may take each session-pairing as a veridical representation of the relative effect of tDCS on these specific occassions, and as such interpret intra-individual differences in how tDCS influences behaviour.

Next, we turned to the correlational structure between the session-pairing psychometric effect, and the EEG activity in the alpha and beta bands. To compress this analysis, we averaged the band-power across the delay epoch. Although this is a concession, in the sense that we previously observed complex band-power changes that did not necessarily translate to a time-averaged directional change with a suitable summary statistic, we stress that this an analysis tool to interrogate how a correlation may be drawn through a common means of abstraction, which necessarily cleaves away some of the complexity we previously observed. *Fig 11* plots, for each participant and psychometric subtraction, the correlation between the mean triple-difference alpha band-power change across the delay epoch, and the z-score of the psychometric difference-in-difference subtraction for each session pairing.

**Figure 11:**
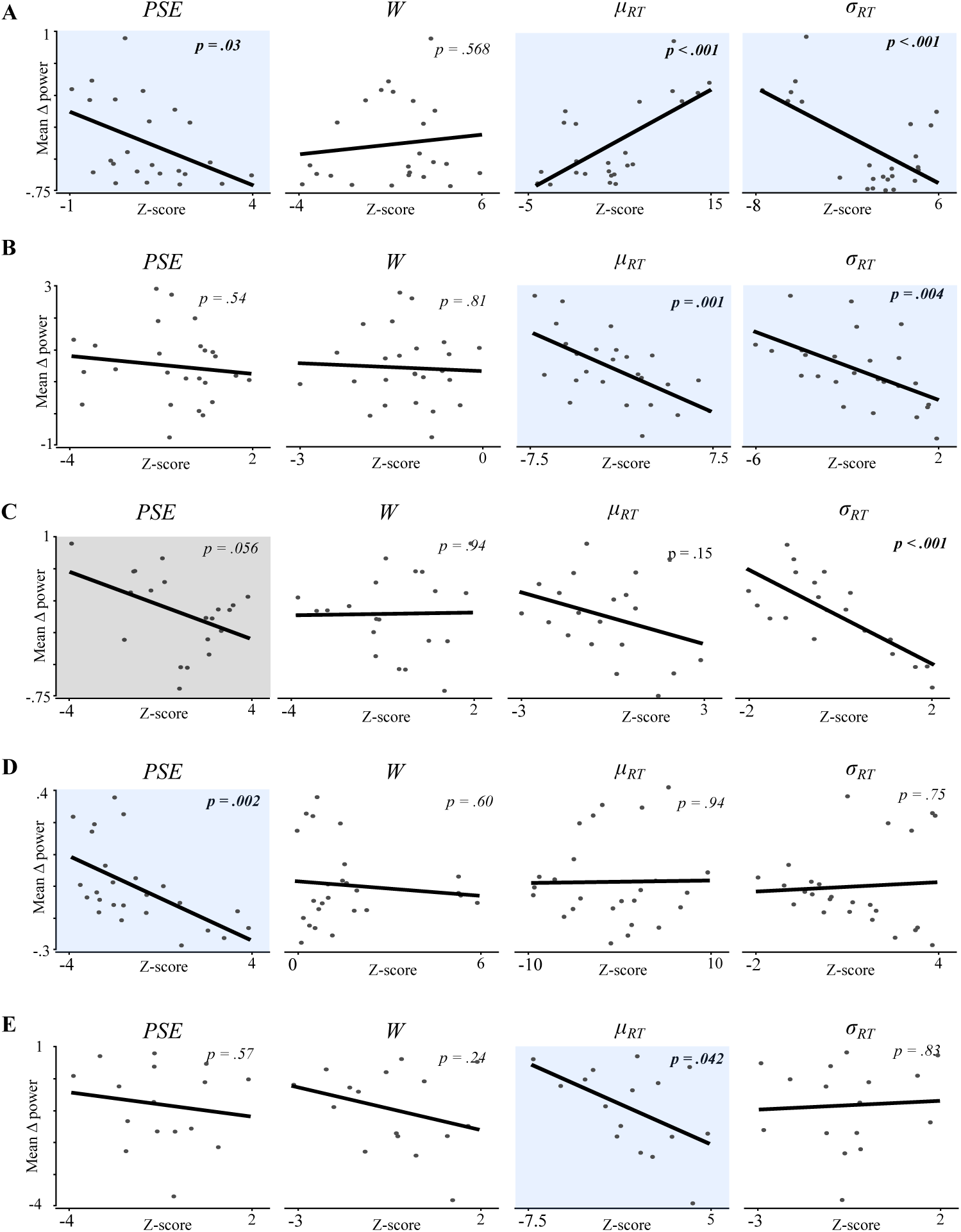
Session-wise correlations between psychoemtrics and EEG beta-band differences. Conventions the same as Fig 10.

To contrast the correlations we report here with the inter-individual level of analysis, we report the consistency between tests at the inter-individual level (where both the psychometric and EEG time-frequency subtractions reached significance), and tests at the intra-individual level (where they significantly correlated). The distinction between what these test is important to state up front. While the former reports whether the these separate significance tests coincided when aggregating all sessions, the latter reports whether there was a relationship between the measures on a session-session basis. For data limitations within-sessions, we neglect to include the metrics of choice history that we previously reported at the inter-individual level. In the inter-individual tests (fig xx), only three non-choice history tests coincided with significance in both the psychometrics and a corresponding change in the reported EEG band-power activity, all with respect to the *PSE*: participant 1, where a increase in the *PSE* subtraction coincided with an early-trial increase in alpha power, participant 3, where a increase in the *PSE* subtraction coincided with an early-trial reduction in beta-power, and participant 5, where a decrease in the *PSE* subtraction corresponded to a reduction in middle-trial alpha power.

Beginning with the alpha-band correlations in *Fig 11*, we see that for both *P*1*_PSE_*_−−_*_>Alpha_, p* = .312 and *P*5*_PSE_*_−−_*_>Alpha_, p* = .7, despite both psychometric and EEG permutations surviving significance when averaged across sessions, we fail to find evidence for a relationship between the psychometric subtractions, as denoted by the z-score on the abscissa, and the corresponding mean alpha power-subtraction. In contrast, we find evidence for several relationships between the psychometric Z-score and the power subtractions 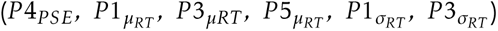 that did not correspond with a coinciding change in both the average psychometric and EEG subtractions. In *Fig 12*, the same analysis is shown for beta-band activity averaged across trials. Here we see that seven significant session-wise correlations 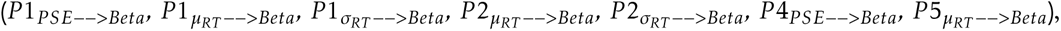 none aligning with the single consistent change in psychometric and EEG beta power at the session-average (*P*3*_PSE_*_−−_*_>Beta_, p* = .056).

**Figure 12:**
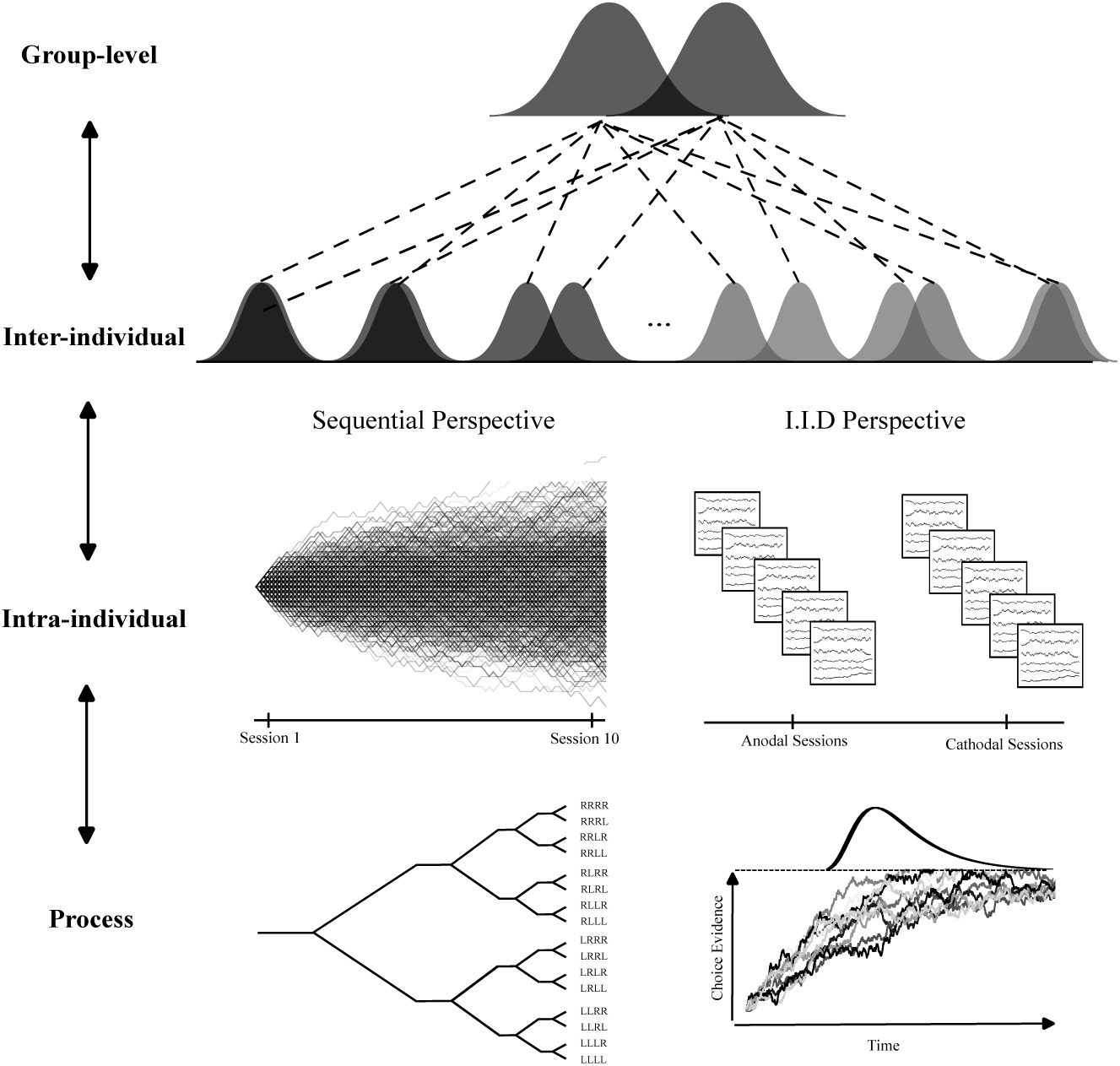
Sequential and I.I.D Perspectives of Statistical Variations in Free Choice Behaviour. Depicted is a hypothetical generative process that supervenes over the correlational structure of free choice saccadic behaviour and EEG activity at the group-level for two intervention conditions (ex: pre-post anodal and pre-post cathodal subtractions). These groups of measurements are composed of a set of individuals each with a set of anodal and cathodal sessions. These sessions in turn can be conceived as independent and identically distributed measurements (I.I.D Perspective), or history-dependent measures arising from a unique sequence (depicted as an ensemble of random walks here, although this choice is arbitrary). For each session measurement, individual trial outcomes can arise from a sequential (depicted as a choice sequence tree with unique historicities) or an ensemble statistical model (a generic bounded integration process here).

## Discussion

In this work, we assessed the influence of HD-transcranial direct current stimulation (HD-tDCS) over the right frontal eye field (rFEF) on free choice behaviour and its neural correlates at multiple levels of abstraction: group-level, inter-individual, and intra-individual. A growing body of evidence has suggested that the exchangeability of results derived from these levels of abstraction are subject to an array of confounds (Horvath et al., 2015; B. Krause and Cohen Kadosh, 2014; Li et al., 2015). Our experimental setup was designed to minimize the influence of inter-individual confounds by individualizing the selection of a brain region involved in the planning and execution of saccadic eye movements with fMRI, using the individual MRIs to guide electrode participant prior to every session, and selecting an electrode montage that minimizes current spread to non-targeted brain regions while maintaining focality (Mikkonen et al., 2020; Villamar et al., 2013). We then developed a multilevel approach to causal inference based on permutation testing of difference-in-differences that allowed us to estimate in a data-driven fashion the probability that changes to psychometrics and EEG time-frequency data were influenced by tDCS. We observed a general discepancy between the statistical relationships uncovered at one level, and the variability that comprised it at a lower level of abstraction, suggesting that tDCS-induced effects may have been different across-sessions.

While our approach may be criticized as a *reductio ad absurdem* with respect to group-level analysis, as the consequences of inter-individual sources of variability are established enough such that blindly combining participants into common statistical units is recognized by many to be flawed (Gallistel, 2012), we argue that the recapitulation of this inconsistency at the intra-individual level of abstraction provides evidence for the interpretation that averaging repeated measures across sessions within-individuals is similarly confounded. Because of the intra-individual variability we report, caution is warranted in interpreting the variability seen in inter-individual studies as evidence of inter-individual variability in tDCS-mediated effects *per se*, in absence of a principled explanation for the intra-individual variability that should be expected under repeated measures of a given experimental context. Without such an explanation, we cannot at this point recommend a single privileged level of analysis (group, inter-individual, intra-individual) at which the effects of tDCS *should* be expected to be suitably generalizable in the context of our task and paradigm, such that repeated measures could reliably separate tDCS-mediated effects from the unknown statistical properties of unmeasured supervening processes.

The early studies evaluating tDCS in humans tested how TMS-induced motor evoked potentials changed over time as a function of the polarity of stimulation (Nitsche and Paulus, 2000, 2001). It was found that anodal stimulation potentiated the size of the motor evoked potential following stimulation, while following cathodal stimulation it was depressed. Although responsible for an influx of human tDCS research, this work has been reevaluated in recent years. Numerous studies have reported substantial inter and intra-subject variability in this paradigm, which may compromise previously reported effects (Chew et al., 2015; Dyke et al., 2016; Horvath et al., 2016; Vergallito et al., 2022; Zappasodi et al., 2018). Some of this variability may be attributed to the specifics of the TMS-MEP protocol (ex Herwig et al., 2001) which are beyond the scope of our work, however other sources of variability are suitably general such that they warrant discussion here.

Most of the narrative surrounding inter-individual variability in response to tDCS concerns differential anatomical and physiological features that may generate individually specific effects subject to masking by group-averaging. With appropriate electrode montages, it is uncontroversial that electric fields can be induced at the cortical surface using tDCS, and this can in turn influence neural activity (Chhatbar et al., 2018; Huang et al., 2017; Opitz et al., 2016; Ruhnau et al., 2018; Vöröslakos et al., 2018; Wiethoff et al., 2014). However, individual differences in anatomical features are substantial enough to predict markedly different current flow at targeted sites, and casts doubt on the anatomical specificity of tDCS (Bikson and Rahman, 2013). This is important for predicting the influence of tDCS on aggregate neural measures, such as EEG, that largely sample activity arising from dipole moments generated by the temporal synchrony of neuronal activity along the columnar architecture of cortical microcircuitry (Buzsáki et al., 2012).

With respect to our paradigm, individual differences in tDCS-induced electric fields can be viewed from at least two vantages. The first is based on predictions derived from a circuit-level reductionist approach (Douglas et al., 1989) to the relationship between neural activity and free choice saccadic behaviour (Heinzle et al., 2007; Soltani et al., 2013). In this view, a dissociation between the predictions of modifying activity in superficial and deep layers of a given brain region may be made, such that the effect of the applied current on the target area may be dependent on not just strength, but also its orientation. (for similar logic see Evans et al., 2022; Rawji et al., 2018, but see Joshi et al., 2023; Laakso et al., 2024). Similarly, variability in the functional definition of the targeted brain region itself may contribute – indeed the frontal eye field in humans is a somewhat nebulous concept that largely inherits etiology from non-human primate studies, and has not converged on a singular definition in humans (Vernet et al., 2014). Accordingly, across our 5 participants there was variability in the localization procedure, as defined by saccadic activity in the functional MRI. Additionally, the variance associated with the induced electric field may be interpreted from a more global perspective with respect to distal neural activity. In our study, the intensity and the variability of the induced current was variable across participants. The degree to which this may cause differential changes to distal activity in the network (either causally through the target region or indirectly through current spread) is challenging to assess, but these unknowns should not be neglected when condensing the results into a mechanistic-reductionist interpretation of how tDCS may influence a single brain region (Bergmann and Hartwigsen, 2021; M. R. Krause et al., 2017; Louviot et al., 2022; Mitchell and Potter, 2024; Polanía et al., 2011; Silberstein and Chemero, 2013).

We show that inter-individual changes to psychometric measures and EEG time-frequency correlates, when averaged across sessions, were composed of a mixture of confirmatory and contradictory results. In some cases, individual session-pairings in our permutation testing analysis largely followed the session-averages, while others session-pairings yielded polarizing effects. All we can say from this definitively is that a single paired session contrasting the effects of anodal and cathodal stimulation are insufficient to provide a representative statistic of the average effect in an individual, as they may diverge substantially from session-averages across repeated measures. However, this intra-individual variability may have arisen from a number of distinct causes that can be roughly divided into two classes. The first would posit that there are intra-individual variables within the causal chain between the measured EEG signal and free choice saccadic behaviour that modify how tDCS impacts our measurements, and that these intra-individual variables change over suitable timescales to confound the independence of sessions. This would essentially propose that i) the causal effect of tDCS on the brain-behaviour relationship is state-dependent, and ii) the underlying states that determine its causal effect are non-stationary across our repeated-measures. Although causal assessments at this level may be challenging (Greene et al., 2023), there is some experimental evidence for state-dependent effects of tDCS at different levels of abstraction – from cellular mechanisms to functional networks (Antal et al., 2007; Paulus and Rothwell, 2016; Vergallito et al., 2023).

A second class of intra-individual causes of our results are agnostic to specific mechanistic predictions of tDCS, but instead are inferential in nature – what should be expected of the statistical structure of variability in absence of a causal effect? And what measures can we deem to be independent? In this manuscript, we developed a permutation-testing approach to assess in a data-driven manner the probability that a change in psychometrics or EEG time-frequency measures were statistically distinguishable by the polarity of the stimulation at three different levels of abstraction. Although this provides us with a useful tool to appraise statistical variations as a function of the intervention we used, the way in which these results are interpreted is far from straightforward, and ultimately depends on the measured processes being independent statistical units (Caie and Blohm, 2024; Mangalam and Kelty-Stephen, 2022; Molenaar, 2004). In the task analyzed in this manuscript, it is fairly straightforward to conceive of different generative models that may confirm or contradict the independence of these statistical units at different levels of abstraction (this division is depicted in *Fig 13*). While our analysis of sequence-dependence at the group and inter-individual level, and prior work on this task, suggest that the outcome of past trials can provide predictive value, and thus provides one counter-example to the issue of using trials as independent measures, we caution against this being used as proof positive of a sequentially-dependent generative process underlying the observations – it would be perhaps trivial to construct a null proposition wherein ensemble models of sequences could be viewed as independent functional units over a longer timescale, such that any history-dependence we observe falls out of an ensemble of sequences that are independent with respect to the timescale of the entire experiment. However, this should not discourage us in discussing potential generative models, if only to clarify how the predictions they generate differ.

The atomization of behavioural measures into discrete functional units is practically axiomatic to the cognitive neuroscience approach (Cisek, 2019; Huk et al., 2018). Under this view, repeated measurements of a certain psychological classification of behaviour are recorded, and variance associated with repeated measures provide evidence for process-level variation that can be used to guide model-based decompositions of behaviour, and help separate expected sources of process-level variance from causal changes in the presence of interventions. Although there is plenty of counterargumentation suggesting that these measurements, and the neural processes they are often correlated with, are not independent – indeed it is challenging to appear algorithmically random (Nickerson, 2002) – the consequences for this on statistical inference are not always made explicit. It is well known in econometrics, a field where the inability to uncover generative models with predictive power is more accepted (and perhaps more obvious), that history-dependence in statistical models produce spurious correlations (Granger and Newbold, 1974; Phillips, 1986). The consequence of this for single paired measurements of history-dependent measures, such as might be expected in the case of our pre-post experiment, is a high rate of false-positive assessments when averaging across trials, and a general difficulty in performing inference of meaningful statistical variations with single session pairings.

Eschewing the atomization of trials is not a death knell for statistical testing in general. While we may have sufficient evidence to cast doubt on the statistical independence of trials in tasks such as ours (despite attempts at ‘randomization’), it may be found that sessions, themselves composed of sequentially-dependent measures, are independent (or *independent enough*) to be used as repeated measurements for inferential studies for a certain class of interventions. Our assessment of intra-individual variability is inconclusive in this regard – while we can say that variance in session-pairings may arise from either tDCS-dependent variance across sessions, or tDCS-independent variability, it is unclear to what degree these contrasting views are separable in a data-driven study like this. What we can conclude is that the sources of variability we observe in free choice saccadic behaviour and its EEG correlates are such that a privileged level of abstraction for data-driven inference of the causal effects of interventions such as tDCS remains elusive. Thus, caution is warranted in using tDCS as a reliable causal intervention in basic science and clinical applications.

## Methods

### Code and Data Availability

Our data is available at https://zenodo.org/uploads/17545450. A public repository to reproduce all analyses is available at https://github.com/BJCaie/tDCS_FEF/.

### Overview

Participants were recruited to undergo an fMRI-guided HD-tDCS study on the role of the frontal eye field in free choice saccade behaviour. Prior to the stimulation sessions, a structural and functional MRI was obtained from each participant in order to localize a region of the fronto-parietal cortex active during saccadic planning and execution. Then, an individualized stimulation setup was designed using 3D models of the neuroanatomy, which in turn guided electrode placement prior to each session. Participants performed 10 sessions of a free choice saccade task over an approximately 5 week period, in a pre/post design alternating between anodal and cathodal stimulation.

### Participants

Five participants took part in the study. All had normal or corrected-to-normal vision. The participants gave written consent prior to undergoing the procedure. The study was conducted with an experimental protocol approved by the Queen’s University General Research Ethics Board, which adheres to the principles of the Canadian Tri-council Policy Statement on Ethical Conduct for Research Involving Humans and the principles of the Declaration of Helsinki (2013).

### Free Choice Task

The participants began by fixating on a central target (*Fig 1A*) projected onto a screen (20” Mitsubishi Diamond Pro [16×12inches] 1280×1024 pixels, 60Hz, contrast 75.2%, brightness 0%). Two targets spaced at 6 degrees visual angle from central fixation in each visual field, were then presented with randomly assigned temporal asynchrony values (0 ms, 16ms, 33ms, 66ms, 99ms) and onset times drawn from a uniform distribution between 750 and 1250ms. The participants were instructed to look at either target, so long as it was done as fast as possible, without anticipating. The participants were informed prior to the beginning of the experiment that the targets would come up at different times relative to each other. The participants were instructed to blink in between trials in order to minimize disruption of stimulus perception. The script for the task was written in MATLAB using Psychtoolbox.

### Behavioural data acquisition and filtering

During each trial, eye movements were measured via an EyeLink 1000 Tower Mount (SR Research, Mississauga, Canada) with a 1000Hz infrared camera that tracked retinal position. Retinal position was calibrated prior to each session. The EyeLink apparatus was placed 60cm from the screen containing the saccade targets. Saccades were calculated offline using a saccade detection algorithm with a velocity criterion of 50°/s, and were individually verified through a custom visual marking program. Trials where the tracker lost the eye (either through tracking error or participant lapse) were excluded. Trials were also excluded if the eye position deviated more than 15 degrees from the horizontal. The first saccade following target onset, or the first saccade within 3 degrees of the future target position prior to target onset, were taken to calculate the saccadic reaction time and endpoint error. Saccades with a positive endpoint horizontal position were given a choice value of 1, and saccades with a negative endpoint horizontal position a choice value of 0.

### fMRI-guided HD-tDCS

To target the site of stimulation, we localized the right frontal eye field using a center-out saccade task during a functional MRI scan. Structural MRI was first acquired on a 3-Tesla Magnetom Trio scanner (Siemens Medical Systems, Erlangen, Germany). Participants rested supine in the scanner and wore a 32-channel head coil. At the beginning of the localizer session, a 176 slice, high resolution T1-weighted anatomic reference volume was acquired using a 3D MP-RAGE sequence (single shot, ascending sequence in the sagittal plane with TR = 1760 ms, TE = 2.2 ms, FoV = 256 mm, flip angle = 9°, and voxel size = 1 mm3). Participants performed center-out saccades, where targets were organized in a ring around a center fixation point. Participants were then tasked to perform saccades from the center point to each target in clockwise succession, pausing for approximately 1 second between each saccade.

Individual structural MRI scans were registered to Talaraich space by manually finding the anterior commisure and translating the cerebrum into AC-PC space. Functional MRI scans taken during the center-out saccade task were collected using a gradient echo planar imaging sequence, and pre-processed using a sinc slice scan time correction interpolation and high-pass filtering before co-registration to the anatomical scan with an initial and fine alignment in Brainvoyager QX. A general linear model was fit to the aligned BOLD activation map to find the voxel most correlated with the saccade period in the vacinity of the pre-central sulcus (Vernet et al., 2014).

HD-tDCS was chosen to maximize current delivery at the target location while minimizing current spread to other brain regions (citation). Briefly, HD-tDCS is a method to improve focalization of current by positioning 4 return electrodes in a concentric ring close to a central electrode. The position of the central electrode was chosen individually for each participant by finding the position on the scalp closest to the voxel chosen as the right FEF using Brainsight (Rogue Research Inc, Montreal Quebec). This mapping was done by performing a 3D reconstruction of the surface of the scalp, and coregistering it to the functional MRI data. A skin reconstruction was performed to identify anatomical landmarks for performing the subject-image registration. Then, a 3D brain reconstruction was overlayed to select a point on the scalp corresponding to the FEF ROI. Prior to each session, the central electrode position on the scalp was then found using a stereotactic neuronavigation system. Participants wore an optical tracking device fixed over the head that was tracked by a position sensory camera located overhead. The position on the scalp corresponding with the target location was then found using an infrared pointer that mapped the position of the pointer to the position on the scalp corresponding to the 3D model .

This scalp position was then recorded with a marker, then a custom HD-tDCS cap was fitted with the central electrode placed over the marked scalp location. The cap was designed so that the central electrode would be surrounded by four pickup electrodes at a radius of 4cm. To lower impedance, hair was dried and separated under the scalp using cotton swabs. Electrodes were 1.5cm diameter Ag/AgCl ring electrodes (UltraStim, Axelgaard Manufacturing, Denmark). Once the center and surround electrodes were placed in the plastic holder, the holder was filled with conducting gel and the stimulation cap was fixed with a chin velcro strap. Continuous direct current was then generated during the stimulation period using a microprocessor-controlled constant current source (neuroConn DC-STIMULATOR MC, Brainbox) . Current was delivered at 2mA for 20 minutes, with a 30 second ramp on and ramp off period.

### Current Modelling

Individualized modelling of the resulting electric field induced by HD-tDCS over the target site was performed in order to predict the strength and orientation of the induced electric field at the target ROI across participants. A tetrahedral head mesh was constructed using the T1-weighted structural MR images using *MRI2Mesh* in SimNIBS (Saturnino et al., 2015). The position of the central electrode was then transformed from BrainVoyager Internal Coordinates to World Coordinates in the SimNIBS GUI. The resulting mesh was used to generate a 4cm ring in the tetrahedral mesh from which the surround electrode positions were selected, then projected on to the scalp in the SimNibs GUI. Electric field simulations were then performed for each subject individually. The central electrode was set to 2mA current, and each surround electrode -.5mA for the anodal current simulation. Cathodal electric fields were taken as the negative sign of the anodal simulations. Electrode parameters in SimNIBS were set to elliptical 1.5cm x 1.5cm, with type “Electrode plus Gel” and electrode thickness of 5mm. Simulation results were visualized using Gmsh 4. Tissue conductivity values were taken from the standard values in SimNibs (Wagner, 2004, Opitz, 2015). A table of the central and surround electrode positions in SimNIBS World Coordinates are shown for each participant in Table 1.

### Session Design

Each individual session was organized into a pre-stimulation period, where EEG and behavioural measures were taken simultaneously, followed by a 20 minute stimulation period where either a net anodal or cathodal current was introduced to the site of the central electrode, and finally a post-stimulation EEG-behavioural recording. In each of the pre/post periods, 5 blocks of 90 trials of the free choice task were recorded, lasting about 20 minutes and totalling 450 trials per session per condition. Sessions were scheduled twice weekly.

### EEG Pre-Processing

EEG data was recorded in the pre and post stimulation periods using the same center-surround electrode setup as for current delivery. EEG data was recorded using a V-Amp 16 channel EEG amplifier (Brain Products, Gilching, Germany) at a sampling frequency of 512 Hz, acquired using a custom protocol in OpenVibe. Reference and ground electrodes were placed on the back of the neck and the right elbow respectively. For each block of trials, EEG channels were independently notch-filtered to remove 60 Hz powerline noise, and then bandpass-filtered between 5 and 30 Hz. The current source density at the central electrode was then estimated using the surface laplacian spatial filter of the central and surround electrodes.

Time-frequency responses of the surface laplacian were computed for each trial individually. EEG data from each trial was aligned to the onset of the fixation cross using a trigger protocol in OpenVibe. The surface laplacian was first detrended to remove slow drifts. Then, a continuous wavelet transform was performed across 25 frequency bands evenly spaced between 5 and 30 Hz. The wavelet transform was performed with a Morlet wavelet of width 5. The average baseline power for the first 50ms of the trial was then subtracted within each frequency band.

### Psychometric Analysis

We quantified choice probability by fitting psychometric functions to behavioral data and extracting parameters corresponding to choice bias (leftward vs rightward) and sensitivity to differences in TOA. For each dataset, the proportion of rightward choices was computed for each temporal onset asynchrony (TOA) bin, and these proportions were fit with a cumulative Gaussian function using the Psignifit toolbox for Python Schütt et al., 2016.

The cumulative Gaussian psychometric function has the general form:

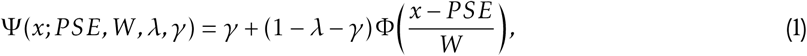

where *x* represents the stimulus value (here, TOA), Φ(·) is the cumulative distribution function (CDF) of the standard normal distribution, *PSE* is the point of subjective equality, *W* determines the slope (and thus sensitivity), *λ* is the lapse rate, and *γ* is the guess rate.

We used the default Psignifit parameterization with the guess rate fixed at *γ* = 0 and the lapse rate set to *λ* = 0.01 (1%) – choices made less than 70ms prior to the onset of the target were excluded from this analysis. From the fitted function, two parameters were extracted for further analysis:

- Point of Equal Selection (PES; *µ*): the TOA at which the probability of a rightward choice is 0.5, quantifying choice bias.
- Width (*W*): the TOA interval corresponding to an increase in rightward choice probability from 5% to 95%, reflecting choice sensitivity.

In addition to choice probabilities, we analyzed reaction times (RTs) for each saccade, defined as the latency between the onset of the choice target and the initiation of the eye movement. For each condition, we computed the mean (*µ_RT_*) and standard deviation (*σ_RT_*) of the RT distribution. The latter was included to take into account the influence of anticipatory or urgency-dependent reaction times that may not change (*µ_RT_*).

### Choice History Analyses

To test whether the inferred effects of transcranial direct current stimulation (tDCS) were history-dependent, we performed additional psychometric analyses conditioned on outcome metrics of the previous trial. Three types of history effects were evaluated.

- Choice Repetition: Trials were grouped based on whether the choice direction (left vs. right) repeated or alternated relative to the previous trial, irrespective of direction.
- Previous Choice Direction: Trials were grouped according to whether the previous choice was to the left or right target, allowing assessment of directional biases.
- Previous Reaction Time: Trials were divided using a median split of the previous trial’s reaction time, contrasting fast versus slow prior responses.

Separate psychometric functions were fit to each subgroup using *psignifit*, and corresponding differences in *µ* and *W* parameters were compared statistically across conditions.

### Multilevel Approach to Causality Testing

We developed a multilevel approach to causality testing that used permutation testing of difference-in-differences to estimate the likelihood of a causal effect of tDCS on psychometrics and EEG activity at multiple levels of abstraction: group-level, individual-level, and session-level. The difference-in-differences approach is a quasi-experimental method used to estimate causal effects of an intervention or treatment when randomized controlled trials are not feasible. Because of this, the approach is most commonly used in the social sciences. Indeed, common approaches to statistical testing in a tDCS study (or a neuroscience study more broadly) would not require this methodology, because it would be assumed *a priori* that a causal effect of tDCS is a population-level phenomenon that can be lawfully uncovered through experimentation with appropriate sample sizes.

However, as we outlined in the introduction, there are numerous reasons to doubt how generalizable the mechanism of action of tDCS may be. Further, this doubt is recapitulated not just at the population-level, due to individual differences that may occur in anatomy and neurophysiology, but also sources of variability that may result in different effects across different sessions (i.e state-dependent effects of tDCS), and changes to outcome metrics within sessions (history-dependence in the underlying processes being measured). Thus, a seemingly randomized experiment (for example, outcome *x* measured pre and post-tDCS) may instead be a series of quasi-experiments with different effects that have been aggregated together. This introduces both the possibility of masking real effects that were dependent on the context of that experimental session, as well as generating spurious findings since the variable controlling the influence of the stimulation is uncontrolled.

In our study, instead of assuming *a priori* the appropriate level at which the data we collected can be considered to be independent measures, we took a multilevel approach in which hypotheses of different classes were evaluated, and the assumptions underlying each were made explicit. First, we collapsed all data across all participants, and used permutation testing of difference-in-differences (detailed below) to quantify the likelihood that the difference between a given behavioural or EEG measure prior to and following anodal stimulation was itself different than the difference prior to and following cathodal stimulation. This allowed us to take into account relative differences in the baseline of different anodal and cathodal sessions (the pre-stimulation period).

Next, we performed this same procedure independently for each participant data. This was possible in our dataset because each participant performed 10 different sessions, allowing for analytical resolution previously impossible in tDCS studies. This involved collapsing all pre-stimulation data on anodal stimulation days together and computing a difference between all post-stimulation data, repeating the same process for cathodal stimulation days, and then taking a final difference-in-difference. The result was a

### Statistical Comparison of Psychometric Parameters

For each psychometric metric *m* ∈ {PES*, W, µ_RT_, σ_RT_* }, we computed a difference-in-difference statistic comparing stimulation polarities and time periods (Pre, Post). Specifically, for each participant and session, we defined:

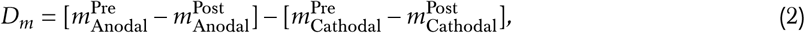

where *D_m_*quantifies the polarity-dependent change in a given metric, controlling for baseline shifts between pre- and post-stimulation conditions.

To obtain the null distribution of *D_m_*, we randomly permuted trial labels across all four condition combinations (*P* = {Anodal, Cathodal}; *C* = {Pre, Post}) while preserving the number of trials per condition. For each permutation *i*, we computed:

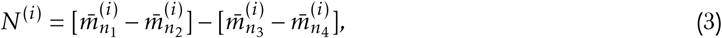

where *n_k_* denotes one of the four resampled condition groups. A total of 500 permutations were generated for each test, providing an empirical estimate of the null distribution under the hypothesis that stimulation polarity had no effect.

The corresponding p-value for each metric was then computed as the proportion of permutations where the permuted statistic exceeded the observed difference-in-difference:

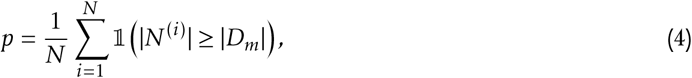

where 1(·) is the indicator function and *N* is the number of permutations. Values of *p <* 0.05 were considered statistically significant.

This procedure was repeated at three hierarchical levels of analysis. First, the group-level, where all sessions were pooled across all participants. Next, the participant-level, where all sessions were combined, but participants were treated separately. Finally, the session-level, where each possible anodal-cathodal session pair was analyzed independently to characterize a distribution of possible polarity-dependent differences across the repeated measurements.

To evaluate whether polarity effects were context-dependent, we extended the same permutation-based framework to a *triple-difference* model by conditioning the psychometric parameters on behavioural context *b* ∈ {previous RT, previous choice direction, repetition}. For each context contrast *b*_1_ versus *b*_2_, the triple-difference statistic was defined as:

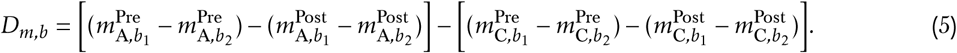

Permutation-based null distributions were again constructed from 500 resamplings of the eight condition–context groups (*P* × *C* × *B* = 8). The resulting empirical p-values quantified the probability that any observed context-dependent polarity effect could have arisen by chance under the null hypothesis of no effect.

### Statistical comparison of EEG Time-Frequency Response

For each participant, session, and stimulation condition, baseline-normalized power spectra were averaged across trials for each of the four primary conditions: stimulation polarity *P* = {Anodal, Cathodal} and stimulation period *C* = {Pre, Post}. The resulting matrices yielded condition-specific power estimates *m_P,C_* (*f, t*) as a function of frequency and time.

To test whether stimulation polarity influenced oscillatory power beyond nonspecific pre/post changes, we computed a difference-in-difference statistic at each time–frequency bin:

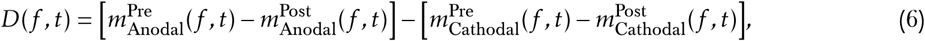

where *D*(*f, t*) quantifies the polarity-dependent modulation of EEG power while controlling for nonstationary baseline shifts between pre- and post-stimulation periods.

To construct the null distribution of *D*(*f, t*), we performed 500 random permutations of trial labels across the four polarity–condition combinations (*P* × *C* = 4), while preserving the number of trials per condition. For each permutation *i*, we computed:

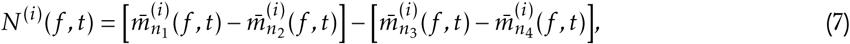

where *n_k_* indexes one of the four permuted condition groups. The resulting permutation distributions provided empirical estimates of the expected variation in *D*(*f, t*) under the null hypothesis of no polarity-specific effect.

Finally, to examine whether EEG effects were modulated by behavioral context, we extended the analysis to a triple-difference model analogous to the behavioral analysis framework. For each behavioral context *b* ∈ {previous RT, previous choice direction, repetition}, we defined:

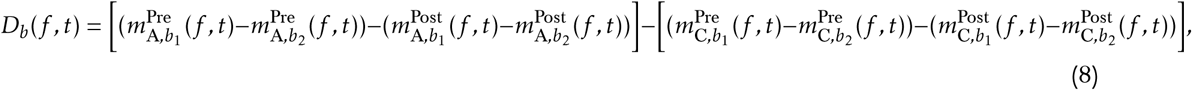

and generated null distributions from 500 permutations across the eight combined condition–context groups (*P* × *C* × *B* = 8).

## Notes

https://github.com/BJCaie/tDCS_FEF

### Competing Interest Statement

The authors have declared no competing interest.

### Summary of Updates

Data and code has been made available, along with minor revisions of the text

### Funding

- Natural Sciences and Engineering Research Council

